# HIV-induced membraneless organelles orchestrate post-nuclear entry steps

**DOI:** 10.1101/2020.11.17.385567

**Authors:** Viviana Scoca, Renaud Morin, Maxence Collard, Jean-Yves Tinevez, Francesca Di Nunzio

## Abstract

HIV integration occurs in chromatin sites that favor the release of high levels of viral progeny, alternatively the virus is also able to discreetly coexist with the host. The viral infection perturbs the cellular environment inducing the remodeling of the nuclear landscape. Indeed, HIV-1 triggers the nuclear clustering of the host factor CPSF6, but the underlying mechanism is poorly understood. Our data indicate that HIV usurps a recently discovered biological phenomenon, called liquid-liquid phase separation (LLPS), to hijack the host cell. We observed CPSF6 clusters as part of HIV-induced membraneless organelles (HIV-1 MLOs) in macrophages, which are one of the main HIV target cells. We describe that HIV-1 MLOs follow phase separation rules and represent functional biomolecular condensates. We highlight HIV-1 MLOs as hubs of nuclear reverse transcription, while the double stranded viral DNA, once formed, rapidly migrates outside these structures. Transcription-competent proviruses localize outside, but near HIV-1 MLOs, in LEDGF-abundant regions, known to be active chromatin sites. Therefore, HIV-1 MLOs orchestrate viral events prior to the integration step and create a favorable environment for the viral replication. This study uncovers single functional host-viral complexes in their nuclear landscape, which is markedly restructured by HIV-1.

## Introduction

Immediately after fusion at the plasma membrane, HIV-1 cores are released into the cytoplasm and move towards the nucleus, while the viral RNA (vRNA) genome begins the process of reverse transcription (RT) into double-stranded viral DNA (ds vDNA) (Blanco-Rodriguez and Di Nunzio 2021; Campbell and Hope 2015; Di Nunzio 2013; V. Scoca and Di Nunzio 2021a). Once imported in the nucleus, the interplay between the HIV-1 DNA genome and the host chromatin compartment is crucial for the fate of the virus-host coexistence. Recent discoveries in early steps of HIV life cycle (Blanco-Rodriguez et al. 2020; Burdick et al. 2020; Dharan et al. 2020; Guedan et al. 2021; C. Li et al. 2021; Rensen et al. 2021; Selyutina et al. 2020; Zila et al. 2021) revise the commonly accepted models of uncoating (loss of the viral capsid) and reverse transcription as exclusively cytoplasmic processes (Farnet and Haseltine 1991; Suzuki and Craigie 2007). It has been found that HIV prompts clusters of the Cleavage and Polyadenylation Specificity Factor Subunit 6 (CPSF6) proteins outside the paraspeckles, but in co-localization with some nuclear speckle factors (Francis et al. 2020; Rensen et al. 2021). However, the nature, the behavior, and the underlying mechanism behind the establishment of these organelles are still unknown.

Here, we distinguish the topological duplicity of the post-nuclear viral entry phases: the early and the late. Early post-nuclear entry steps involve the formation of the HIV– induced membraneless organelles (HIV-1 MLOs), composed of CPSF6 clusters enlarged with viral factors. Late post-nuclear entry steps are characterized by the completion of the double-stranded vDNA synthesis and integration. The final products of the reverse transcription process are recruited into the pre-integration complex (PIC) eventually for viral integration and transcription. The vDNA nuclear dynamics and its surrounding chromatin landscape dictate the evolution of HIV infection (Maldarelli 2016; M. E. Sharkey et al. 2000; M. Sharkey et al. 2011). However, the mechanisms behind the fate of the proviral DNA in the nucleus remain unclear (Liu et al. 2020; Olson et al. 2019). These are challenging to study, also due to the limits imposed by available technologies. Real-time imaging approaches can provide new insight into unprecedented spatial information in single cells by tracking individual viruses.

Here, we perform single cell studies. We describe early post-nuclear entry steps in which the viral infection enhances the phase separation of CPSF6, building HIV-1 MLOs. Our results emphasize post nuclear entry steps as finely regulated by the interplay between viral and host components. Of note, CPSF6 contains intrinsically disordered mixed-charge domains which are responsible for the formation of liquid condensates *in vitro* (Greig et al. 2020) and they may account for the formation of HIV-1 MLOs in infected cells. Membraneless organelles are established through a recently discovered biological phenomenon called liquid-liquid phase separation (LLPS) in which molecules transition from a liquid state to a more solid state, similarly to oil and water demixing (Alberti et al. 2019; Banani et al. 2016; Brangwynne et al. 2009). Our data reveal that HIV-1 MLOs undergo biomolecular condensates’ rules, concentrating viral and host factors in the same niche and hosting nuclear reverse transcription.

For the study of the late post-nuclear entry steps, we labelled the newly synthesized double-stranded (ds) vDNA through the HIV-1 ANCHOR system (Blanco-Rodriguez et al. 2020). The ds vDNA is the only form included in the mature PIC able to give rise to a functional infection. Through the tracking of the vDNA and transcriptional vRNA we found that transcriptional viral foci are excluded but remain in the vicinity of HIV-1 MLOs. Our finds suggest HIV-1 MLOs’ role in creating a surrounding microenvironment favorable for viral replication. We provide new insights into how HIV reprograms and markedly restructures the nuclear environment to orchestrate viral replication steps.

## Results

### HIV-1 MLOs are hubs of viral reverse transcription

Upon nuclear entry, the virus reprograms the nuclear localization of the host Cleavage and Polyadenylation Specificity Factor Subunit 6 (CPSF6) from being randomly dispersed in the nucleoplasm to discrete clusters (Figure 1A) (Bejarano et al. 2019; Francis et al. 2020; Rensen et al. 2021). It has been suggested that the nuclear entry of the viral capsid that interacts with CPSF6 (Buffone et al. 2018; Price et al. 2014) could drive the formation of CPSF6 clusters (Francis et al. 2020). Indeed, viruses carrying the capsid mutation N74D, unable to bind CPSF6, do not prompt CPSF6 clustering formation (Supplementary Figure S1A) (Selyutina et al. 2020). On the other hand, this phenomenon could also be prompted by the overexpression of CPSF6 (Chaudhuri et al. 2020) due to the viral infection.

**Figure 1.**
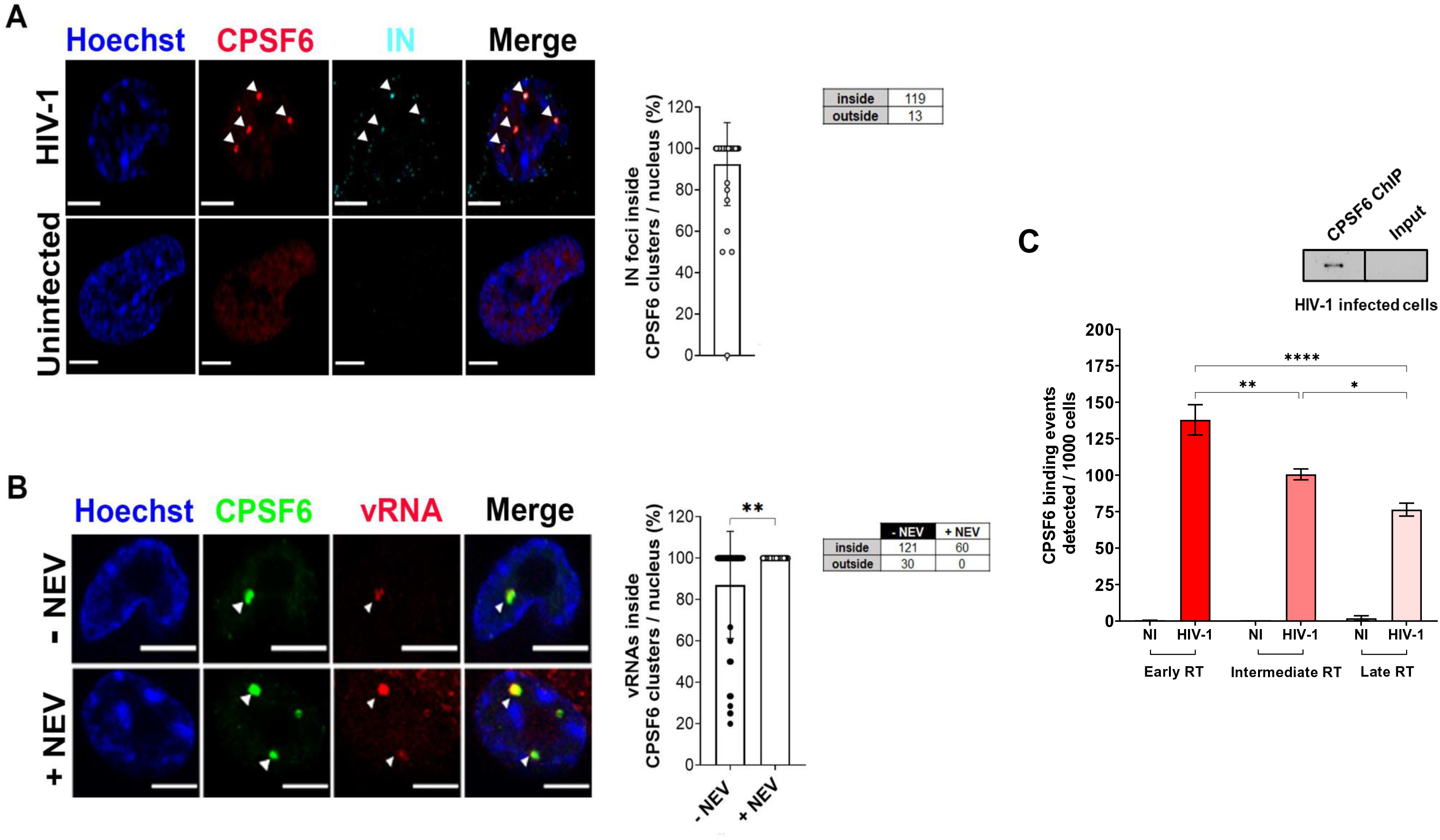
CPSF6 clusters host HIV-1 viral complexes and are hubs of nuclear reverse transcription. **A)** Confocal microscopy image of HIV-1 infected THP-1 cells (MOI 10, 30 h p.i.), compared to uninfected cells. The graph shows the percentages of IN associated with CPSF6, per nucleus (40 cells, 132 IN foci) ± SD. Three independent experiments were performed. **B)** Confocal microscopy images of Immuno-RNA FISH in infected THP-1 (MOI 20, 2 days p.i.) treated or not with NEV (10 µM). The graph shows the percentages of vRNA foci associated with CPSF6 per cells, per condition (cells: 56 (-NEV), 34 (+NEV), vRNA: 151 (-NEV), 61 (+NEV)) ± SD. Unpaired t test, **=p ≤ 0.01. Two independent experiments were performed. **C)** Chromatin Immunoprecipitation of endogenous CPSF6 coupled to real-time PCR of THP-1 cells infected with HIV-1 (MOI 5), 2 days p.i. Different reverse transcription (RT) products were amplified normalized on the input ± SD. One-way ANOVA followed by Tukey’s multiple comparison test, *=p ≤ 0.05, **=p ≤ 0.01, ****=p ≤ 0.0001. On the right, Western blot of CPSF6 of ChIP products in infected cells compared to the input. Scale bar: 5 µm.

Here, we characterize the HIV-1 post-nuclear step and its interplay with the host nuclear landscape in human macrophages. First, we evaluated the nuclear location of the viral endogenous integrase (IN) protein by the immuno-labelling of the hemagglutinin (HA) tag fused to the C-terminal domain of the IN (Blanco-Rodriguez et al. 2020; Petit et al. 2000). We confirmed that the majority of the viral IN proteins accumulate in foci occupied by CPSF6 clusters (Figure 1A and Supplementary Figure S2A) (Francis et al. 2020; Muller et al. 2021; Rensen et al. 2021). To better define the relevance of these viral-host clusters, we performed an immuno-RNA smiFISH (single molecule inexpensive FISH), specific for HIV-1, and we uncovered the presence of incoming viral RNA (vRNA) inside CPSF6 clusters (Figure 1B). Upon the block of reverse transcription through Nevirapine (NEV) treatment, we observed incoming vRNA genomes 100% located inside CPSF6 clusters. Likewise, in physiological infection conditions at 48 hours (h) post-infection, without the drug treatment, vRNA foci can be detected inside CPSF6 clusters (Figure 1B) in most of the cells. Only few cells showed a mixed population of vRNA foci inside and outside, suggesting that the detected vRNAs can have a different nature (Figure 1B). Thus, we asked the functional role of the co-existence between the incoming vRNA and IN proteins sequestered inside CPSF6 clusters. We performed in infected and uninfected macrophage-like cells (THP-1), chromatin immunoprecipitation (ChIP) of the endogenous CPSF6 protein followed by real-time PCR (Figure 1C and Supplementary Figure S1B). Early, intermediate, and late reverse transcription (RT) products were amplified using appropriate primers. Thus, different-stage RT products are associated with CPSF6, suggesting an RT activity inside CPSF6 clusters, (Figure 1C). Our results corroborate previous data showing EdU-labeled viral DNAs (vDNAs) clustering in nuclear niches in macrophages (Francis et al. 2020; Rensen et al. 2021), where nuclear RT activity has been detected once an inhibitor of the reverse transcriptase, the Nevirapine, was removed (Rensen et al. 2021). Taken together, our new data combined with previous results, indicate that CPSF6 clusters serve as hubs of nuclear reverse transcription. We called these hubs hosting viral and host factors: HIV-1 membraneless organelles (HIV-1 MLOs).

### HIV-1 MLOs are phase separated condensates

Next, we investigated the phenomenon underlying the establishment of HIV-1 MLOs. Membraneless organelles (MLOs) allow the fine regulation of complex functions in a spatiotemporal manner (Banani et al. 2016; Fare et al. 2021). MLOs are generated by a newly characterized liquid-liquid phase separation (LLPS) process, that can be driven by the spontaneous assembly of intrinsically disordered regions (IDRs) present in some proteins (P. Li et al. 2012). Of note, CPSF6 contains disordered regions, such as arginine-enriched mixed-charge domains that have been evoked as responsible for the phase separation features of this protein (Greig et al. 2020). To investigate whether LLPS aids the virus to navigate into the nuclear space through CPSF6, we performed the following experiments. First, we observed an increase in CPSF6 intensity exclusively in the nucleus of infected cells (Figure 2A), in line with a recent observation that HIV-1 induces an increase of CPSF6 proteins (Chaudhuri et al. 2020). This observation suggests that CPSF6 during viral infection undergoes to LLPS, which is a concentration-dependent phenomenon (Alberti et al. 2019). We also observed a reduction of CPSF6 clusters in infected cells treated with 1,6 Hexanediol (1,6 HD). This drug dissolves protein droplets driven by IDRs, commonly used to test LLPS *in vitro*. However, these findings should be viewed with caution. Indeed, our experiment also shows toxicity (nuclear morphology is drastically affected) due to this compound (Supplementary Figure S2H), as previously reported (Wheeler et al. 2016). Second, we found that CPSF6 clusters increase twice their volume along the time post-infection (24 h p.i. vs. 72 h p.i.) and that they are spherical (Figure 2B), typical shape of liquid droplets (Brangwynne et al. 2011; Strom and Brangwynne 2019). The mean intensity of CPSF6 signal positively correlates with the mean intensity of IN proteins in the cluster (Pearson’s r= 0.8563) (Supplementary Figure S2A). Third, to evaluate their dynamics, we live-tracked HIV-1 MLOs. We used differentiated THP-1 cells stably expressing CPSF6 fused to mNeonGreen (Zhong et al. 2021). Of note, in about all cases the endogenous-only CPSF6 clusters were occupied by the exogenous fluorescent proteins and CPSF6 mNeonGreen clusters carried viral components, such as the integrase (IN) (Supplementary Figure S2B). Thus, we performed time-lapse epifluorescence microscopy at 5-min intervals to capture fusion events (Figure 2C and Supplementary Video S1). We could appreciate the high dynamism of the HIV-1 MLOs, which keep restoring their circular shape (Figure 2C and Supplementary Figure S2C). Along our videos acquired between 24 to 72 h post infection, HIV-1 MLOs constantly evolved and performed fusion and fission (Figure 2B and C, Supplementary Figure S2C-E), which characterize LLPS-condensates. Next, to evaluate whether HIV-1 MLOs are characterized by high mobility of the proteins essentially depending on diffusion, we performed fluorescence recovery after photobleaching (FRAP). CPSF6 free proteins in the nucleoplasm show high diffusion (Supplementary Figure S2F). In the context of infected cells, we observed that after the laser irradiation (200 µs/px) the florescence of CPSF6 cluster started to recover at ∼ 2 minutes, indicating the exchange of fluorescent CPSF6 molecules inside and outside the MLO (Figure 2D and Supplementary Video S2), contrary to the viral IN cluster which does not recover after bleaching (Supplementary Figure S2G). Nevertheless, this experiment is suggestive of LLPS behaviour, which is in line with other fluorescent recoveries detected for other factors that follow LLPS properties in nuclear speckles (Hong et al. 2020; Marzahn et al. 2016; Taylor et al. 2019) (Supplementary Figure S2I). We have also observed that CPSF6 clusters fully colocalize with the marker of nuclear speckles (NSs), SC35 (Figure 2E), in line with previous studies (Francis et al. 2020; Rensen et al. 2021). CPSF6 IDRs are the major players to promote NS residence (Greig et al. 2020). We remarked that CPSF6 and SC35 signals mainly occupy inter-chromatin space with low-Hoechst stain (Figure 2E). We also confirmed that SC35 shows rapid recover of the fluorescence signal after photobleaching (Supplementary Figure S2I) (Marzahn et al. 2016).

**Figure 2.**
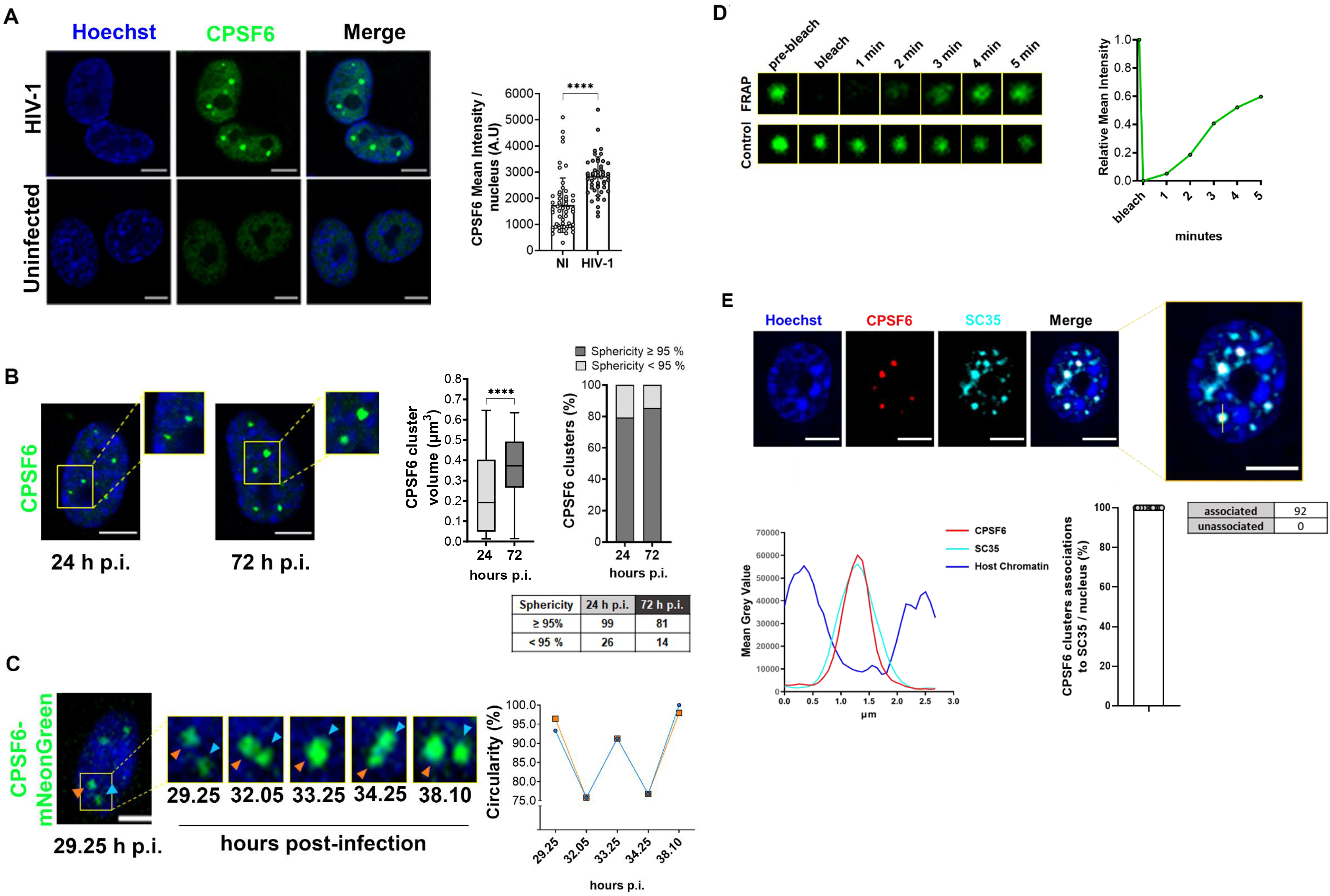
Phase separation properties of HIV-1 MLOs. **A)** Confocal microscopy images of THP-1 cells infected with HIV-1 (MOI 5, 24 h p.i.), compared to uninfected cells. On the right, mean intensity of CPSF6 per nucleus ± SD (n cells = 49, 55). Unpaired t test, ****=p ≤ 0.0001. Two independent experiments were performed. **B)** Confocal microscopy images of THP-1 cells infected with HIV-1 (MOI 5) + NEV (10 µM) comparing 24 h and 72 h p.i. On the right, box plots of the volume (median: ∼ 0.19 vs ∼ 0.37 µm^3^) and percentage of CPSF6 clusters that show ≥ or < 95% sphericity value (n = 125, 95). Unpaired t test, ****=p ≤ 0.0001, ns = not-significant. 3D analysis of two independent experiments. **C)** Frames extracted from a 10 hours-time-lapse microscopy in 2D (Supplementary Video S1) of THP-1 cells expressing CPSF6 mNeonGreen infected with HIV-1 (MOI 10). The graph shows the circularity (normalised ratio between the contour and interior expressed as a percentage) along the memebraneless organelles fusion-fission event. Representative of 6 independent experiments. **D)** Frames extracted form a Fluorescence Recovery After Photobleaching (FRAP) time-lapse (Supplementary Video S2) in THP-1 cells expressing CPSF6 mNeonGreen infected with HIV-1 (MOI 10, 3 days p.i.). Representative of 3 independent experiments. The graph shows the recovery of the signal curve. Pre-bleach signal is set to 1 and bleach signal is set to 0. **E)** Confocal microscopy image of THP-1 cell infected with HIV-1 (MOI 10, 3 days p.i.). The graph on the left shows the intensity profile of Hoechst, CPSF6, SC35 signal along the segment crossing the MLO (yellow). On the right, the percentage of CPSF6 colocalizing with SC35 per nucleus ± SD, 3D analysis of two independent experiments. Scale bar: 5 µm.

Our results inscribe HIV-1 MLOs in the plethora of biomolecular condensates (Alberti et al. 2019). Our observations suggest that HIV-1 MLOs assemble through a multistep process initiated by CPSF6-virus complexes then assembling via phase-separation. The condensates seem to exhibit a central stable structure decorated by highly dynamic CPSF6 proteins that exchange with the surrounding environment.

### The newly synthesized double-stranded viral DNA separates from HIV-1 MLOs

To decipher the nuclear location of the newly synthesized ds vDNA we benefited from the HIV-1 ANCHOR DNA tagging system (Supplementary Figure S3A) (Blanco-Rodriguez et al. 2020). Once fully reverse transcribed, the HIV-1 genome carrying the bacterial sequence ANCH3 is specifically recognized by OR-GFP protein, which is expressed in the target cells, genetically modified by the lentiviral vector (LV) OR-GFP (Blanco-Rodriguez et al. 2020) (Supplementary Figure S3A). The accumulation of OR-GFP, which is a modified version of the bacterial ParB protein (Graham et al. 2014; Sanchez et al. 2015), on the ANCH3 sequence generates bright nuclear signals (Figure 3A and B, Supplementary Figure S3 B and C). OR protein binds exclusively the ds DNA (Saad et al. 2014). We cloned ANCH3 in the nef gene, thus, HIV-1 ANCHOR system exclusively detects the final products of the RT, because the ANCH3 sequence is one of the last sequences to be converted into ds DNA (Supplementary Figure S3A). Thus, the system tracks the vDNA forms that will potentially be engaged in the PIC. Unlike previous studies based on viral protein tracking (Burdick et al. 2017; Burdick et al. 2020; Chin et al. 2015; Francis and Melikyan 2018; Francis et al. 2020; Lelek et al. 2012), we are now able to directly visualize in fixed and live cells individual nuclear vDNA (GFP spots). The high specificity of HIV-1 ANCHOR system to detect the newly synthesized vDNA is shown by the correlation between the nuclear GFP puncta and the MOI (multiplicity of infection) used (Supplementary Figure S3B). In addition, the RT inhibitor, NEV, completely prevents the detection of vDNA (Supplementary Figure S3B). We also assessed that HIV-1 ANCHOR can detect both vDNAs, episomal and integrated forms. Indeed, this labeling system can reveal intranuclear GFP spots in infected cells treated with an inhibitor of integration, Raltegravir (RAL), or by using an integration defective virus (HIV-1 ANCH3 IN_D116A_) as confirmed by qPCR (Supplementary Figure S3C). Importantly, we also observed that HIV-1 ANCHOR is sufficiently sensitive to allow the detection of a single provirus, as shown by the results of HIV integrations (ALU PCR) and of single cell imaging of a selected clone (Supplementary Figure S3D).

**Figure 3.**
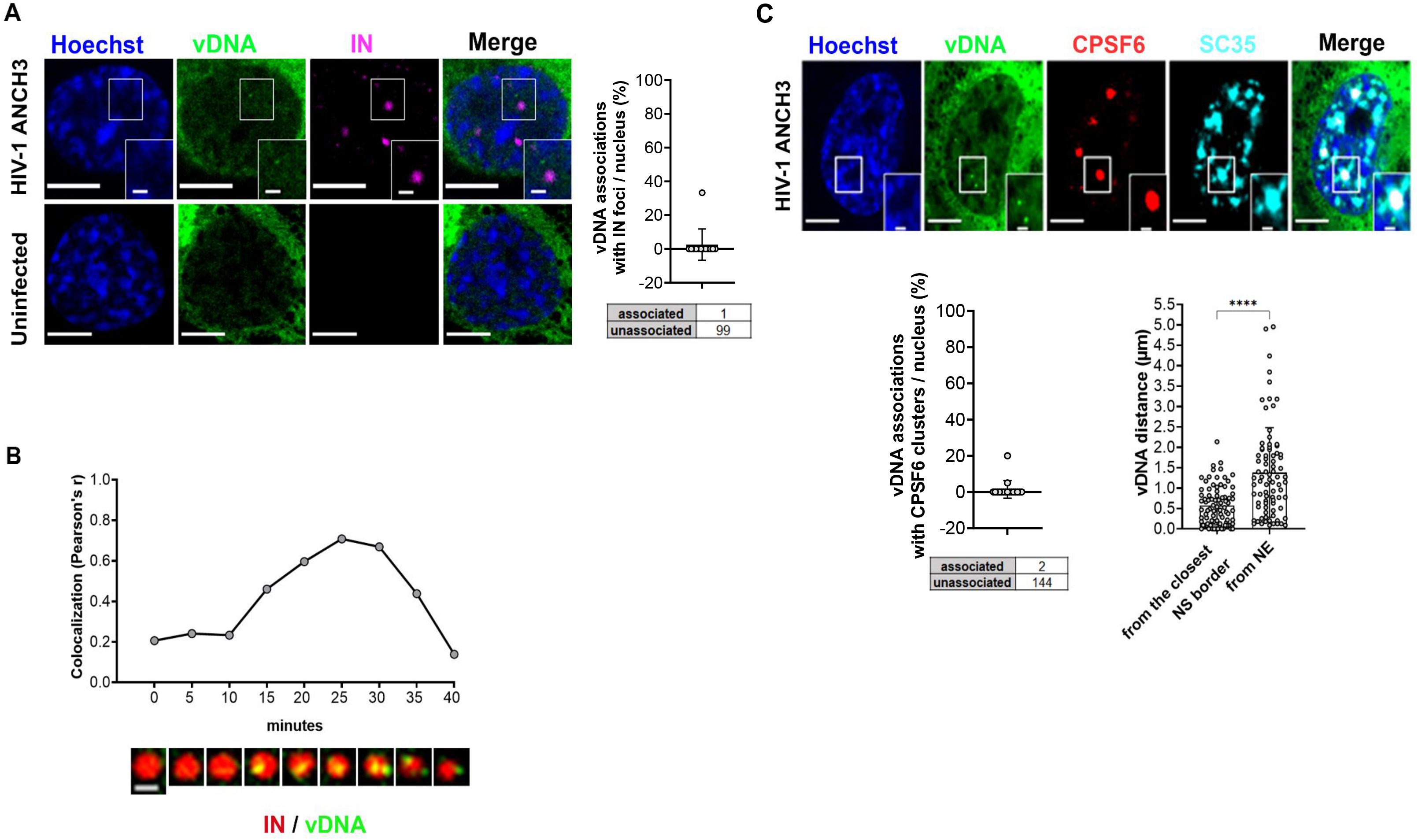
Complete reverse transcribed products are released from HIV-1 MLOs. **A)** Confocal images of THP-1 cell expressing OR-GFP, infected with HIV-1 ANCH3 (MOI 10, 30 h p.i.) compared to uninfected cell. The graph shows the percentages of vDNAs associated with IN, per nucleus (13 cells, 100 vDNAs) ± SD, 3D analysis of two independent experiments. **B)** Cropped frames from a time-lapse microscopy (Supplementary Video S3) of vDNA (green) and IN (red) dynamics in THP-1 cells (HIV-1 ANCH3 GIR, 80 h p.i.). Pearson’s correlation of the two signals per 2D frame along the time. Representative of 3 independent experiments. Scale bar: 1 µm. **C)** Confocal image of THP-1 cells expressing OR-GFP infected with HIV-1 ANCH3 (MOI 10, 3 days p.i.). On the bottom, the graphs display: the percentage of vDNA associated with CPSF6 clusters, per nucleus (17 cells, 146 vDNAs) and the distance ± SD of each vDNA from the closest nuclear envelope (NE) point and SC35-marked nuclear speckle border (7 cells, 83 vDNAs), 3D analysis of two biological replicates. Unpaired t test, ****=p ≤ 0.0001. Scale bar: 5 µm

Intriguingly, when we labelled late reverse transcripts with respect to the IN foci hosted in HIV-1 MLOs, we found a complete separation, since their association was extremely rare (Figure 3A). The HIV-1 IN-vDNA nuclear dynamics represent a long-standing and debated topic. There are, indeed, multiple challenges due to the short-time nature of the event or the technical limitations of vDNA and IN visualization. Thus, to confirm our data we performed live-imaging, asking whether we could pinpoint the dynamic evolution of this separation, coupling the HIV-1 ANCHOR system with the Gag-integrase-Ruby (GIR) virus (generous gift from E. Campbell) (Dharan et al. 2016; Hulme et al. 2015) (Figure 3B and Supplementary Video S3). The virus is produced using a viral genome carrying ANCH3 sequence together with the GIR plasmid, incorporating fluorescent IN proteins in the viral particle. Performing multiple videos in THP-1 cells during a non-synchronized infection we found IN foci (marking HIV-1 MLOs) in which several vDNA puncta arose before their separation (Figure 3B and Supplementary Video S3). We analysed the association of the two signals that rapidly decreases (Figure 3B). According to these data, there is formation of final products of reverse transcription inside HIV-1 MLOs. Of note, in fixed cells we can label the nascent vDNAs via EdU incorporation, that remain mostly associated with the CPSF6 clusters, as opposed to the late reverse transcription products which are mainly detected outside the CPSF6 clusters (Supplementary Figure S3E). The vDNA/IN separation data in fixed and live cells indicated that final products of the RT are released from HIV-1 MLO sites once synthesized, in line with a recent study (Muller et al. 2021). At three days p.i., nearly all double-stranded late reverse transcripts were excluded from HIV-1 MLO cores (Figure 3C), as also suggested from the separation of the vDNA from the IN puncta (Figure 3A B). The majority of the viral IN visible by confocal remain sequestered in CPSF6 clusters (Figure 1A), likely part of the multiple viruses retained in these organelles. Overall, our results indicate that IN proteins that are part of individual HIV PIC can be hardly visualized. The limit of resolution of a conventional light microscope can account for the inability to track a single HIV-PIC, even if IN proteins form high-order structures in an individual PIC (Ballandras-Colas et al. 2017; Hare et al. 2009; Passos et al. 2017; Passos et al. 2020).

Next, we investigated the nuclear space distribution of the final products of the reverse transcription. We computed the 3D distances of the ds vDNAs from the boundary of the closest NS or nuclear envelope (NE). We calculated that the vDNAs are located closer to NS than to the NE, at a distance ∼ 0.57 µm in average (Figure 3C). Therefore, our results show that the vDNAs labelled by HIV-1 ANCHOR are mainly excluded from HIV-1 MLOs.

### HIV-1 active proviruses locate outside HIV-1 MLOs

Taking in consideration that the ds vDNAs are found outside HIV-1 MLO (Figure 3A-C), we wondered about the location of viral transcription sites that allow the viral replication. Thus, we coupled HIV-1 ANCHOR with RNA FISH to co-detect the vDNA and the vRNA. The vDNA/vRNA association analysis highlights the transcription sites of the virus in both differentiated THP-1 cells and primary monocyte-derived macrophages (MDMs) (Figure 4A). Not all vDNAs were associated with a transcriptional focus, because the episomal forms are less actively transcribed than integrated vDNAs. However, some vDNAs co-localize with bright vRNA spots (RNA FISH), suggesting that these are active proviruses.

**Figure 4.**
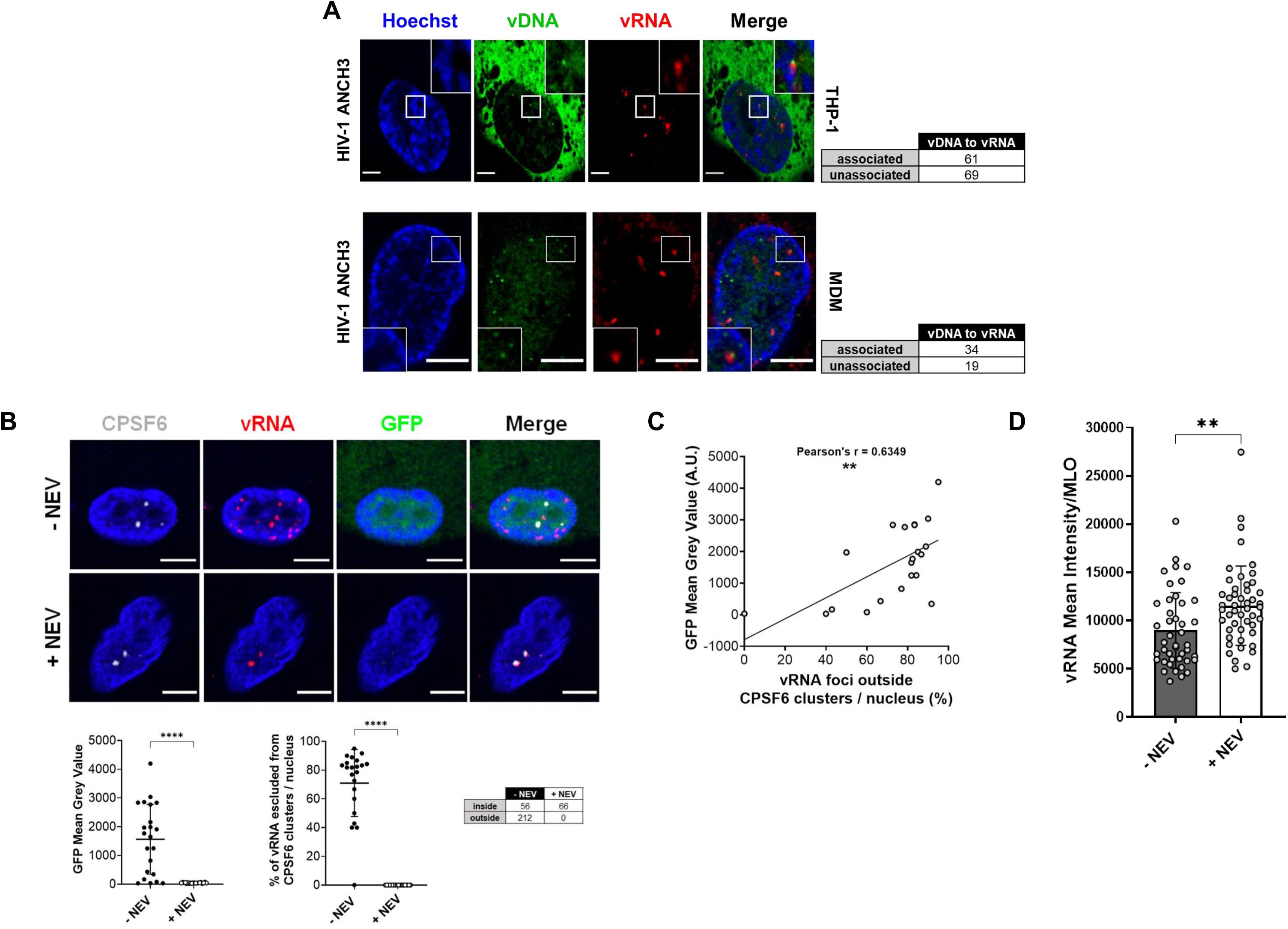
Active proviruses are excluded from HIV-1 MLOs. **A)** Confocal images of THP-1 and primary macrophages (MDMs) cells expressing OR-GFP, infected with HIV-1 ANCH3 (THP-1: MOI 20, 3 days p.i., MDMs: MOI 40, 4 days p.i) 3D analysis. **B)** Confocal images of Immuno-RNA FISH in primary MDMs infected with HIV-1 GFP (MOI 20) ± NEV (10 µM), 2-3 days p.i. The scatter plots show the GFP intensity ± SD per cell and the percentage of vRNA outside CPSF6 clusters ± SD per nucleus (cells: 22 (-NEV), 20 (+NEV); vRNAs: 268 (-NEV), 66 (+NEV)); Unpaired t test, ****=p ≤ 0.0001. **C)** Correlation in infected cells (Pearson’s r coefficient, **=p ≤ 0.01) of GFP intensity and % of vRNA foci exclusion from CPSF6 clusters (22 cells). **D)** vRNA intensity ± SD per CPSF6 clusters in ± NEV conditions; unpaired t test, **=p ≤ 0.01. All the analyses were performed in 3D from two independent experiments. Scale bar: 5 µm.

Next, we focused on the nuclear location of active proviruses in MDMs infected with HIV-1 GFP reporter virus to monitor viral expression (Figure 4B and C, Supplementary Figure S4A). Through the immuno-RNA FISH we were able to co-visualize CPSF6 clusters and HIV-1 vRNA as incoming genomes and viral transcription foci. We observed that at three days p.i., MDMs treated with NEV contain only incoming vRNAs sequestered in CPSF6 clusters (Figure 4B), similarly to THP-1 cells (Figure 1B). On the other hand, untreated cells presented a mixed population of vRNA foci inside and outside HIV-1 MLOs (Figure 4B). The percentage of excluded vRNA foci correlates with GFP expression (Pearson’s r ∼0.63) suggesting that these foci are actively transcribing proviruses (Figure 4B and C).

Moreover, we found that the mean intensity of the vRNA signal in HIV-1 MLOs (MLOs have been selected for the sphericity >95% (Supplementary Figure S4B) was higher in average in NEV treated cells compared to untreated cells (Figure 4D), likely due to an ongoing RT process (Figure 1C). The MLOs-excluded vRNA foci can be also tracked down by coupling HIV-1 ANCHOR (vDNA detection) to immune-RNA FISH (vRNA detection), indicating that the sites of viral transcription locate outside HIV-1 MLOs (Supplementary Figure S4C). These results are supported by the decrease in number of excluded foci in infected cells treated with RAL (Supplementary Figure S4D) which do not show HIV-1 GFP reporter signal and consistent with the notion that episomal forms transcribe much less than proviruses (Dupont et al. 2021).

The immune-RNA FISH highlights both vRNA foci inside and outside the MLO (Figure 5A). On the contrary, when we applied MCP-MS2 system (kind gift from E. Bertrand) (Tantale et al. 2016), we were able to exclusively visualize vRNAs located outside HIV-1 MLOs. Upon block of viral integration (RAL), we could rarely detect vRNA foci, due to the low level of transcription of episomal forms (Dupont et al. 2021), while in absence of drug we could highlight several vRNA foci, indicating that these bright spots are active proviruses located outside CPSF6 clusters (Supplementary Figure S5A). Thus, MCP-MS2 applied in the context of HIV, exclusively detects viral transcription foci (Figure 5B and Supplementary Figure S5A). Besides, we observed that MCP-GFP is hindered in binding incoming vRNA and it is filtered out by HIV-1 MLOs. In fact, upon NEV treatment no viral RNA species were detected inside CPSF6 clusters (Figure 5B). The specificity of MCP-MS2 in the binding vRNA transcription foci could be due to the steric hindrance from the capsid that shields the vRNA genome (C. Li et al. 2021; Muller et al. 2021; Zila et al. 2021) or because MLOs can act as selective filters (Alberti et al. 2019). However, further investigations are needed to better characterize this phenomenon. In addition, this approach allows the live tracking of individual proviral transcription foci by coupling it to HIV-1 ANCHOR technology (Supplementary Figure S5B). Overall, MCP-MS2 is a valuable tool to study the interplay between active proviruses and the surrounding nuclear landscape. Thus, we used MCP-MS2 system to compute the 3D distances of viral transcription foci from the closest NS boundary or the NE. We observed that viral replication sites locate in the NS-surrounding chromatin (avg ∼ 0.57 µm) (Figure 5C). These results, uploaded on bioRxiv in 2020 (Viviana Scoca et al. 2020) have been confirmed by a recent study from another group (Burdick et al. 2022). It has been found by genomic approaches that HIV-1 favours integration in speckle associated domains (SPADs) (Francis et al. 2020), known to be active chromatin loci (Kim et al. 2020). Now, our study allows to spatially distinguish and visualize individual vRNA transcription foci in single cell. We were able to appreciate the nuclear topology of active proviruses.

**Figure 5.**
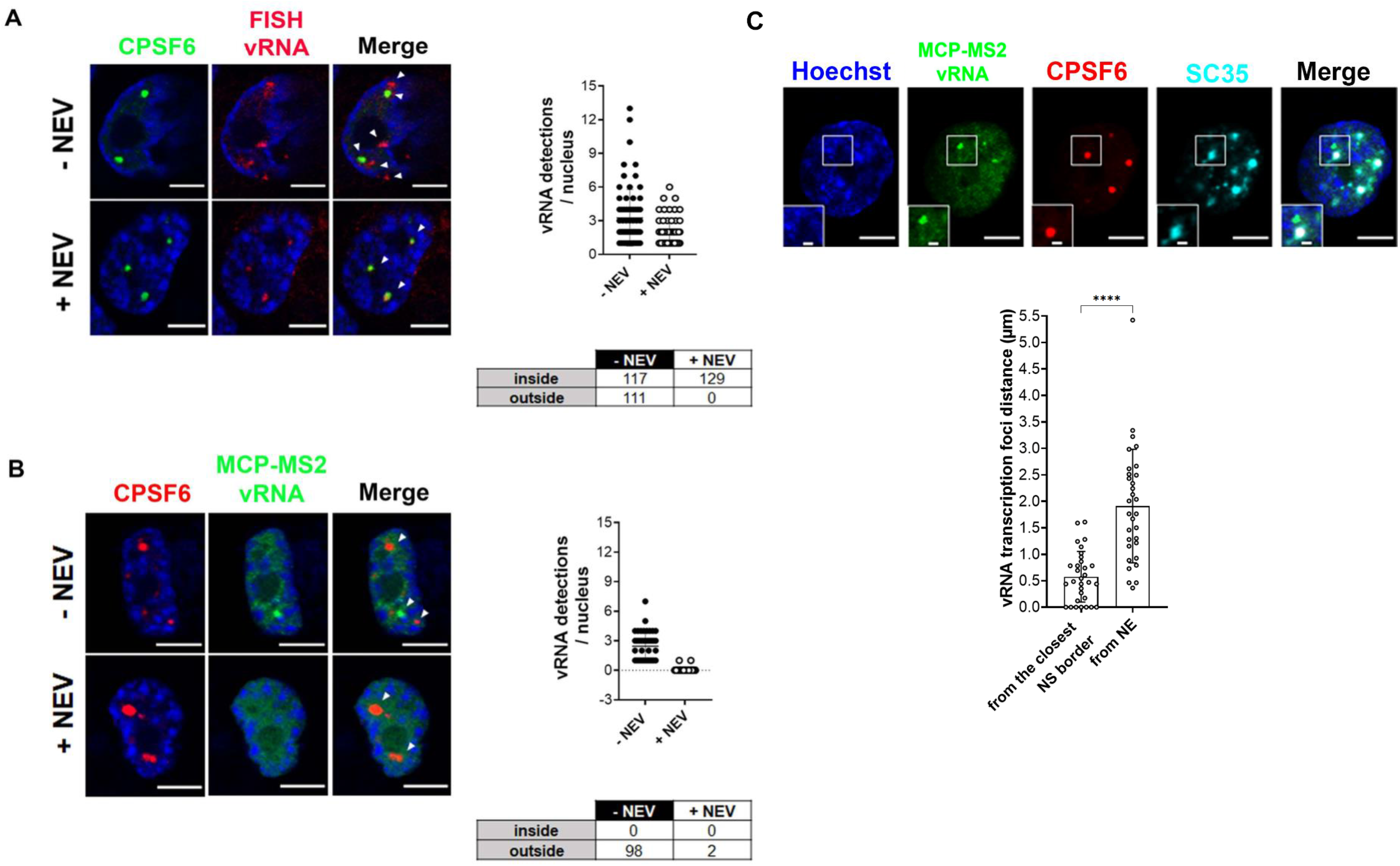
MCP-MS2 RNA labelling detects transcription foci only. **A)** Confocal images of Immuno-RNA FISH in THP-1 cells infected with HIV-1 ± NEV (10 µM), (MOI 20, 3 days p.i.). The graph shows the number of vRNA per nucleus, the table shows the amount of vRNA inside or outside CPSF6 clusters (n = 70 cells (-NEV), 68 cells (+NEV)). Two independent experiments were performed. **B)** Confocal images of THP-1 cells expressing MCP-GFP infected with HIV-1 ANCH3 MS2 ± NEV (10µM), (MOI 2.5, 3 days p.i.). The graph shows the number of vRNA per nucleus, the table shows the amount of vRNA inside or outside CPSF6 clusters (n = 40 cells (-NEV), 44 cells (+NEV)). Three independent experiments were performed. **C)** Confocal images of THP-1 cell expressing MCP-GFP infected with HIV-1 ANCH3 MS2 (MOI 2.5, 3 days p.i.). The graph shows the distance of each RNA focus from the closest nuclear envelope (NE) point and SC35-marked nuclear speckle border (31 vRNAs, 8 cells) ± SD, 3D analysis of two independent experiments. Unpaired t test, ****=p ≤ 0.0001. Scale bar: 5 µm, inset: 1 µm.

### The HIV-1 replication occurs in LEDGF-chromatin sites near HIV-1-MLOs in macrophage-like cells

We explored the nuclear landscape surrounding HIV-1 MLOs using HIV-1 ANCHOR and MCP-MS2 labelling systems. Lens Epithelium-Derived Growth Factor (LEDGF) (Ciuffi et al. 2005), which is a host partner of the viral IN (Cherepanov et al. 2003; Ciuffi et al. 2005; Emiliani et al. 2005; Ferris et al. 2010; Hare et al. 2009; Lelek et al. 2015; Llano et al. 2006; Shun et al. 2007; Shun et al. 2008; H. Wang et al. 2014), participates in HIV-1 integration sites distribution. We observed that LEDGF forms clusters independently of viral infection in macrophage-like cells (THP-1) (Supplementary Figure S6). LEDGF is located in active chromatin regions because its PWWP domain has been shown to interact with H3K36me3 (Pradeepa et al. 2012; Pradeepa et al. 2014), which is a marker of euchromatin. This post-translational modification is enriched in HIV-1 integration sites (Ciuffi et al. 2005; Lelek et al. 2015; Vansant et al. 2020; G. P. Wang et al. 2009). In this study we found LEDGF to be associated with the proviruses forming a complex visible by imaging (Figure 6A), dissociated from CPSF6 clusters (Figure 6A). In average ∼64% of vDNA and ∼73% of vRNA foci per nucleus were found in complex with LEDGF (Figure 6A). LEDGF is also known to interact with splicing factors (Singh et al. 2015). This agrees with our data showing vDNA and vRNA localized in average at ∼ 0.57 µm from the closest NS (Figure 3C and 5C). Our data support HIV-1-targeted active chromatin surrounding HIV-1 MLOs. More in detail, the foci of viral transcription locate at less than 1 µm from the closest HIV-1 MLO (CPSF6+SC35), whereas the total vDNAs (episomal or integrated forms) are more distant, in average ∼1.8 µm (Figure 6B).

**Figure 6.**
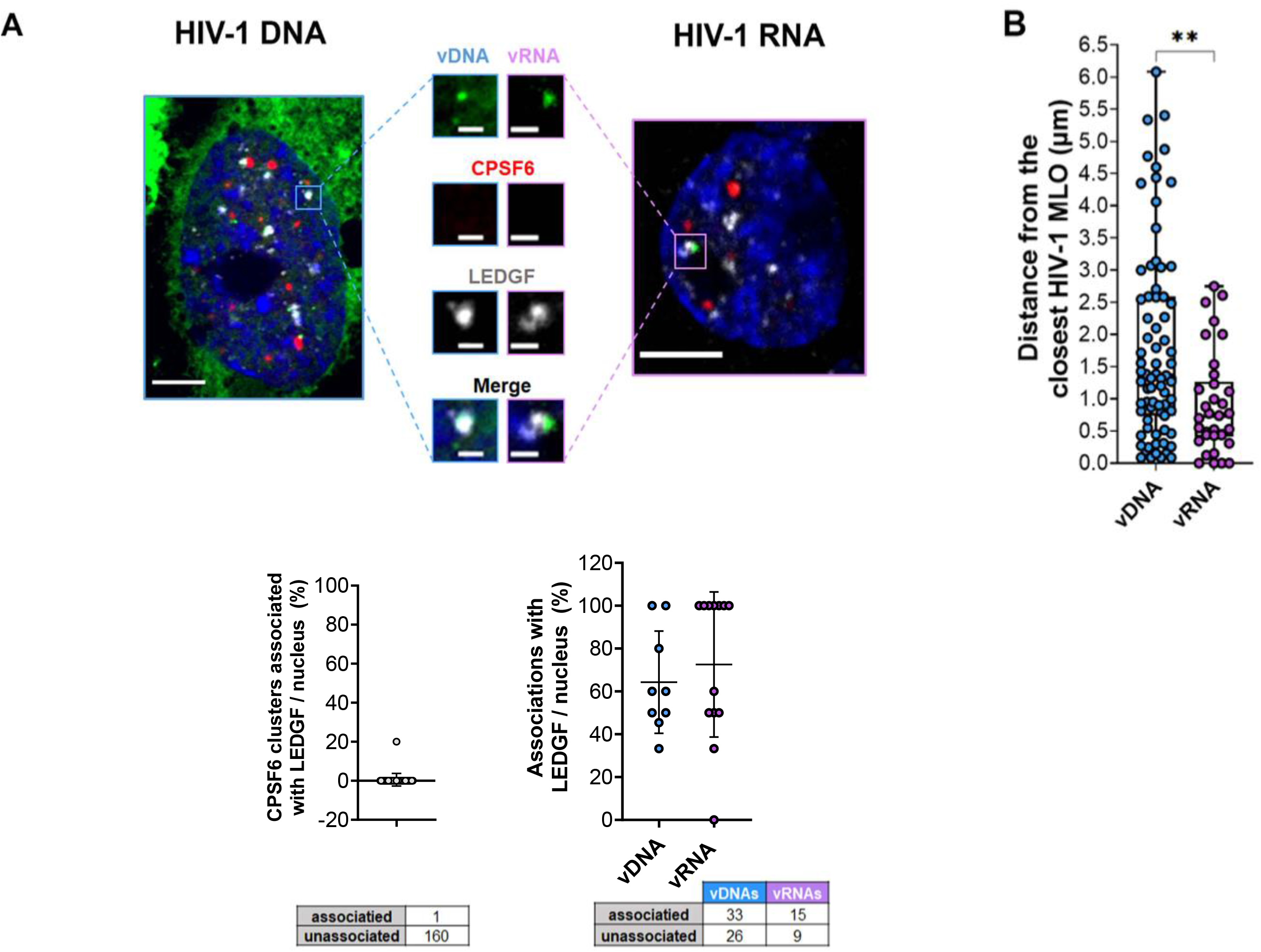
HIV-1 active proviruses locate in LEDGF-marked chromatin neighboring HIV-1 MLOs. **A)** Confocal image of THP-1 cells expressing OR-GFP infected with HIV-1 ANCH3 (MOI 10, 3 days p.i.) and expressing MCP-GFP infected with HIV-1 ANCH3 MS2 (MOI 2.5, 3 days p.i.). The plot shows the percentage of CPSF6 clusters associated with LEDGF per nucleus (38 cells, 161 CPSF6) ± SD and the percentage of vDNA or vRNA associated with LEDGF cluster per nucleus ± SD. Two independent experiments were performed. **B)** Scatter plot with bars indicating the distance of vDNA or vRNA from the border of the closest (SC35+CPSF6) MLO ± SD (n = 75 vDNAs, 7 cells; 34 vRNAs, 8 cells). Unpaired t test, **=p ≤ 0.01, 3D analysis of two biological replicates. Scale bar: 5 µm, inset: 1 µm.

Overall, our results indicate that HIV-1 MLO-neighboring active chromatin regions, marked by LEDGF and in proximity of NS, favour HIV-1 integration, creating a suitable environment for viral replication.

## Discussion

Pathogens, such as HIV-1, able to integrate their genome into the host DNA have evolved to target chromatin both to favor the release of new progeny and to optimize their coexistence with the host (Bushman 2003). Even though HIV-1 infection is characterized by the silent persistence of the virus in the host cell, HIV needs to rapidly generate high levels of virions during the acute phase, intuitively explaining the preference of active genes as integration targets (Ciuffi et al. 2005; Di Nunzio et al. 2013; Lelek et al. 2015; Maldarelli 2016; Schroder et al. 2002).

Thus far, CPSF6 has been described as crucial partner of HIV-1 capsid (Buffone et al. 2018; Lee et al. 2010; Price et al. 2014) and as player in HIV-1 integration site selection (Achuthan et al. 2018; Buffone et al. 2018; Sowd et al. 2016). When CPSF6 was first discovered, a deletion mutant of CPSF6 unable to translocate in the nucleus was found as a potent inhibitor of HIV-1 infection (Lee et al. 2010). Here, we provide new insights on the properties of CPSF6 protein clusters induced by the infection, and we unravel their unexpected role in HIV-1 pre-integration steps in non-dividing cells, known to be an important cellular reservoir (Ganor et al. 2019; Kruize and Kootstra 2019; Real and Bomsel 2019; Veenhuis et al. 2021). CPSF6 proteins have been previously shown to cluster in NSs upon viral infection (Francis et al. 2020; Rensen et al. 2021). We observed that CPSF6 proteins form condensates that occupy the interchromatin space (Figure 2E) and host multiple viral components, at early times post-infection (Figure 1A-B). These host-viral condensates form even in the absence of reverse transcription (Figure 1B), implying their relevance in early post-nuclear entry stages. Multiple viral EdU-labelled DNA copies strongly colocalize with CPSF6 clusters (Supplementary Figure S3E) (Francis et al. 2020; Rensen et al. 2021). Indeed, the ChIP of the endogenous CPSF6, followed by real time PCR in infected THP-1 cells (48 h p.i.), shows an abundance of early, intermediate, and late reverse transcription products. We also observed that the amount of the vRNA inside CPSF6 clusters decreases if there is an RT in progress compared vRNA sequestered in CPSF6 clusters in presence of NEV (Figure 4D). These data are in line with our previous study that highlighted the existence of a nuclear reverse transcription, observed when the NEV was removed, restoring the reverse transcription inside CPSF6 clusters (Rensen et al. 2021).

We named these nuclear viral/host niches HIV-induced membraneless organelles “HIV-1 MLOs” (Viviana Scoca and Di Nunzio 2021b) and we investigated their liquid-liquid phase separation (LLPS) properties in the cell. CPSF6 contains intrinsically disordered mixed-charge domains which are responsible for the formation of CPSF6 condensates *in vitro* (Greig et al. 2020). On the other hand, it seems that the binding of CPSF6 to the viral capsid is essential to observe CPSF6 clusters (Supplementary Figure S1A). Thus, we propose that CPSF6 interacts with viral complexes via the CA binding domain and the phase separation properties aid the virus to navigate inside the nuclear space to form HIV-1 MLOs. We show that HIV-1 MLOs match LLPS properties in the cells, being dynamic functional condensates and evolving along infection (Figure 2). Furthermore, the recovery of CPSF6 fluorescence after photobleaching indicates the importance of CPSF6 equilibrium maintenance in these structures (Figure 2D). CPSF6 phase-separation with viral reverse transcription complexes may be induced as response to the nucleoplasm perturbation by HIV, to find a new protein(s) balance. Of note, CPSF6 plays key biological functions, such as in the mRNA processing. We observed an increase of the amount of this protein upon viral infection (Figure 2A) and this phenomenon could ensure a bimodal role of CPSF6, one for the biology of the host cell and the other for the viral replication.

Our data hinted that HIV-1 MLOs play a role in pre-integration steps. To characterize the later nuclear stages of the virus, we exploited HIV-1 ANCHOR system, which is the only DNA labelling approach permitting in fixed and live cells to track HIV-1 ds DNA, including proviruses (Supplementary Figure S3D, Figure 4A and Supplementary Figure S5B). The final reverse transcription products rarely co-localize with IN foci, leaving HIV-1 MLO (Figure 3A-C). This may be due to the inability of confocal microscopy to detect the INs that are part of the PIC or because the integration event occurs at the surface of the HIV-1 MLO to rapidly go further. We can live-track both, proviral and episomal forms using HIV-1 ANCHOR but we cannot distinguish them, however we can follow the moment of their release from HIV-1 MLOs. At three days post infection, in non-dividing cells, the vDNA population is composed of both episomal and proviral forms. Next, results obtained by the confocal microscope indicate that the vDNAs labeled by HIV-1 ANCHOR are excluded from the MLOs and the majority locates around 0.57 µm from the closest SC35-marked nuclear speckle (Figure 3C). Likely, HIV exploits the induced CPSF6 clusters to generate pre-integration complexes (PICs), which then will depart from these interchromatin sites (Figure 3B). Notably, HIV-1 ANCHOR is sensitive enough to enlighten single proviruses (Supplementary Figure S3D) and through the coupling with virus-specific RNA smiFISH (Tsanov et al. 2016) it is possible to reveal transcriptionally active proviruses (Figure 4A). The RNA FISH in primary macrophages displays viral transcription foci excluded from CPSF6 clusters, as previously hinted by the vDNA location. The vRNA foci excluded from MLOs positively correlate with the GFP expression used as viral reporter gene (Figure 4B and C), indicating that the viral replication occurs outside HIV-1 MLOs, while the vRNA genomes reverse transcribe inside (Figure 1C, 3B and 4D).

We finally focused on the nuclear landscape of viral expression, exploiting MCP-MS2 tagged viruses, which mark viral transcription foci only, contrary to the RNA FISH (Figure 5B and Supplementary Figure S5A and B). Viral RNA transcription foci locate outside HIV-1 MLOs, but in proximity of NSs (Figure 5C). LEDGF associates with highly-spliced transcriptional units through its direct interaction with several splicing factors (Singh et al. 2015) and determines HIV-1 integration sites distribution in active chromatin sites (Ciuffi et al. 2005; Ferris et al. 2010; Shun et al. 2007). However, a complex formed by the vDNA or vRNA and LEDGF was never highlighted by fluorescence imaging (Figure 6A). We uncovered that most of the cells present a high percentage of vRNA foci associated with LEDGF (Figure 6A), highlighting the active nature of HIV-1 targeted domains. Lastly, the vRNA transcription foci are closer to CPSF6 + SC35 condensates compared to the total ds vDNA nuclear forms, suggesting that the HIV-1 MLOs-neighbouring regions are preferred hubs of viral replication (Figure 6B).

Taken together our results show that CPSF6 phase-separates to generate speckles-enlarged MLOs prompted by the virus, likely, to usurp functions linked to speckle factors. We propose the following model: HIV-1 MLOs host viral complexes and their nuclear reverse transcription, but once the ds vDNA is synthesized, this is released from them. The nuclear speckle-associated chromatin is a favorable environment for viral replication since active proviruses locate at less than 1 µm from the closest HIV-1 MLOs (Figure 6B). Our study supports how new single-cell level approaches are pivotal in the study of functional nuclear dynamics of viruses, to discriminate the cells susceptible to fuel viremia or that concur to viral persistence.

## Materials and Methods

### Cell lines, primary cells and generation of HIV-1 ANCH3 single-cell clones

THP-1 (ATCC) are immortalized monocytic cells that once seeded, were differentiated into macrophage-like cells under Phorbol 12-myristate 13-acetate (PMA) treatment (160 nM). HEK 293T cells (ATCC) are human embryonic kidney cells used to produce lentiviral vectors and HIV-1. HeLa P4R5 are β-galactosidase reporter cells expressing CD4, CXCR4, CCR5 (Charneau et al. 1994). HeLa MCP-GFP cells stably express MCP-GFP bacterial fusion protein (Tantale et al. 2016). THP-1 cells were cultivated in RPMI 1640 medium supplemented with 10% Fetal Bovine Serum (FBS) and 1% penicillin-streptomycin solution (100 U/mL). HEK293T and HeLa P4R5 cells were cultivated in DMEM medium supplemented with 10% Fetal Bovine Serum (FBS) and 1% penicillin-streptomycin (100 U/mL). Primary monocyte-derived macrophages (MDMs) were obtained by PBMCs buffy coat isolation from healthy donors’ blood via EFS (Etablissement Français du Sang, Paris), through density gradient centrifugation with Ficoll 400. The PBMCs were then incubated at 37°C for 2 hours, next, non-adherent cells were washed away, and complete RPMI 1640 medium was supplemented with M-CSF (10 ng/mL). After 3 days, the medium was changed with RPMI without M-CSF and cells were left to differentiate for another 4 days before proceeding with experiments.

HIV-1 ANCH3 single-cell clone was generated starting from HeLa P4R5 cells which were infected with HIV-1 ANCH3 (MOI 1). Cells were diluted to 1 cell per well, in 96-well plates. Cell-clone colonies were tested for β-galactosidase expression to check viral infectivity (kit Roche #11758241001). Positive clones were transduced with the lentiviral vector (LV) CMV-OR-GFP for the imaging of HIV-1 ANCH3 provirus. All cells were kept in incubator at 37°C and 5% CO_2_.

### Plasmids, lentiviral vectors, and viral productions

HIV-1ΔEnvIN_HA_ΔNef plasmid encodes for the ΔEnvHIV-1 LAI (BRU) viral genome where the integrase (IN) protein is fused to the hemagglutinin (HA) tag (Petit et al. 1999, 2000). HIV-1ΔEnvIN_HA_ΔNef ANCH3 (HIV-1 ANCH3) was obtained through ANCH3 insertion: ANCH3 sequence was cloned by PCR using the template plasmid pANCH3 as we previously described in Blanco et al. (Blanco-Rodriguez et al. 2020). The ANCHOR^TM^ technology and sequences are exclusive property of NeoVirTech (Germier et al. 2017; Mariame et al. 2018). HIV-1ΔEnv IN_HA_(D116A) ΔNef ANCH3 (HIV-1 ANCH3 IN_D116A_) was obtained by insertional mutagenesis using the QuikChange II XL Site-Directed Mutagenesis kit (Agilent #200522), for integrase (IN) mutation at D116. The lentiviral vector plasmid pSICO-CPSF6-mNeonGreen encoding CPSF6 fluorescent fusion protein was a gift from Zandrea Ambrose (Addgene plasmid # 167585 ; http://n2t.net/addgene:167585 ; RRID:Addgene_167585).

The lentiviral vector SC35-mRuby plasmid was kindly provided to us by R. Parker (University of Colorado). HIV-1ΔEnvIN_HA_ΔNef ANCH3 MS2 plasmid was obtained by inserting the MS2×64 sequence, from pMK123-MS2×64 plasmid, (kindly provided by E. Bertand, IGH, Montpellier) (Tantale et al. 2016) in HIV-1ΔEnvIN_HA_ΔNef ANCH3. HIV-1 ΔEnv Δnef IRES GFP plasmid encodes for the ΔEnvHIV-1 NL4.3 GFP viral genome (Rensen et al. 2021). The lentiviral vector plasmids CMV OR-GFP and CMV OR-SANTAKA were obtained by cloning OR-GFP/OR-SANTAKA cassette (plamids from NeoVirTech) in a pTripCMV (ΔU3) plasmid through PCR using restriction site, AgeI and SgrDI.

Lentiviral vectors and HIV-1 viruses were produced by transient transfection of HEK293T cells through calcium chloride coprecipitation. Co-transfection was performed as follows: for lentiviral vectors 10 µg of transfer vector, 10 µg of packaging plasmid (gag-pol-tat-rev) and 2.5 µg of pHCMV-VSV-G envelope plasmid; for VSV-G HIV-1ΔEnv viruses: 10 µg HIV-1ΔEnv plasmid, 2.5 ug of pHCMV-VSV-G plasmid.

Lentiviral vectors and viruses for transduction/infection of THP-1 and MDM cells were produced in combination with 3 µg of SIV_MAC_ Vpx (Durand et al. 2013). HIV-1 ANCH3 IN_D116A_ has been also produced in combination with GIR (Gag-IN-Ruby plasmid) (Dharan et al. 2016; Hulme et al. 2015) for IN/vDNA live microscopy. After the collection of the supernatant 48h post-transfection, lentiviral particles were concentrated by ultracentrifugation for 1 h at 22000 rpm at 4°C and stored at −80°C. Lentiviral vectors and viruses were titred by qPCR in HEK293T cells, 3 days post transduction.

### Chromatin Immunoprecipitation with qPCR

Four hundred million of THP-1 cells infected or not with HIV-1 (MOI 5) were fixed with 1% formaldehyde for 15 min and quenched with 0.125 M glycine, then sent to Active Motif Services (Carlsbad, CA) to be processed for ChIP-qPCR. In brief, chromatin was isolated by the addition of lysis buffer, followed by disruption with a Dounce homogenizer. Lysates were sonicated and the DNA sheared to an average length of 300-500 bp. Genomic DNA (Input) was prepared by treating aliquots of chromatin with RNase, proteinase K and heat for de-crosslinking, followed by ethanol precipitation. Pellets were resuspended and the resulting DNA was quantified on a Nanodrop spectrophotometer. Extrapolation to the original chromatin volume allowed quantitation of the total chromatin yield. An aliquot of chromatin (40 µg) was precleared with protein A agarose beads (Invitrogen). Genomic DNA regions of interest were isolated using 4 µg of antibody against CPSF6 (#NBP1-85676). Complexes were washed, eluted from the beads with SDS buffer, and subjected to RNase and proteinase K treatment. Crosslinks were reversed by incubation overnight at 65°C, and ChIP DNA was purified by phenol-chloroform extraction and ethanol precipitation. Quantitative PCR reactions were carried out in triplicate on specific genomic regions using SYBR Green SuperMix (BioRad). The resulting signals were normalized for primer efficiency by carrying out qPCR for each primer pair using Input DNA. Different RT products were amplified using the following primers, early RT products: 5′-GCCTCAATAAAGCTTGCCTTGA-3′ and 5′-TGACTAAAAGGGTCTGAGGGATCT-3′; intermediate RT products: 5′ - CTAGAACGATTCGCAGTTAATCCT-3′ and 5′ -CTAT CCTTTGAT GCACACAATAGAG-3′; late RT products: 5′-TGTGTGCCCGTCTGTTGTGT-3′ and 5′-GAGTCCTGCGTCGAGAGAGC-3’. A group of control genes were also amplified to show the enrichment of HIV-1 specific DNA in the infected condition. De-crosslinked ChIP products were checked by western blot against CPSF6 (same ChIP antibody (#NBP1-85676)), loading 37.5 ng of DNA per sample.

### RNA FISH and IMMUNO-RNA FISH

THP-1 cells were seeded on coverslips (12mm #1, ThermoFisher) and differentiated with PMA (160 nM) for 48h. For RNA FISH, the cells were transduced with OR-GFP LV (MOI 0.5) and then infected with HIV-1 ANCH3 (MOI 20) for 3 days. The medium was always supplemented with PMA. Primary MDMs were seeded before differentiation on coverslips and after 7 days, transduced with OR-GFP LV (MOI 10) and infected with HIV-1 ANCH3 (MOI 40) for 4 days. The day of fixation the cells were washed with PBS and fixed with 4% PFA for 15 minutes and incubated in 70% ethanol at −20°C at least for one overnight. Primary smiFISH probes have been designed against HIV-1 *pol* sequence and containing a shared readout sequence for secondary probe alignment. Twenty-four smiFISH probes (Supplementary Table S1) against HIV pol were designed with Oligostan (Tsanov et al. 2016) and purchased from Integrated DNA Technologies (IDT). Primary probes were pre-hybridized with a secondary FLAP probe conjugated to Cy5 fluorophore (Eurofins) through pairing with the readout sequence. Washes and hybridization were performed with Stellaris Buffers (WASH buffer A, WASH buffer B, Hybridization Buffer; LGC Biosearch Technologies), following the manufacturer protocol. Hybridization with the probe was carried out at 37°C in a dark humid chamber for 5 hours.

For the immuno-RNA FISH THP-1 cells were seeded on coverslips and differentiated with PMA (160 nM) for 48h and then infected with HIV-1 (MOI and time points indicated in figure legends). NEVIRAPINE (NEV) 10 µM or RALTEGRAVIR (RAL) 20 µM were used to block respectively DNA synthesis or integration. Primary MDMs were seeded before differentiation on coverslips and after 7 days infected with HIV-1 GFP reporter virus for 2-3 days.

Fixation was carried on in 4% PFA for 15 minutes and then the coverslips where incubated with a blocking/permeabilization solution (PBS-BSA 1%, 2mM Vanadyl ribonucleoside complexes (VRCs) solution (Sigma #94742), TRITON-X 0.3%) for 1 h. Then, the coverslips were incubated with anti-CPSF6 1:400 (#NBP1-85676) in blocking/permeabilization solution for 1h, washed and incubated for 45 min with anti-Rabbit secondary antibody, at room temperature, in the dark, in humid chamber. The coverslips were fixed again in PFA 4% for 10 min. Subsequent RNA FISH and mounting protocol was carried out as mentioned before, but the hybridization with the probes was carried on overnight.

Finally, all the coverslips were stained with Hoechst 33342 1:5000 (Invitrogen #H3570) for 5 minutes and mounted on glass slides (Star Frost) with Prolong Diamond Antifade Mountant (Life Technologies #P36970). Confocal microscopy was carried out with a Zeiss inverted LSM700 microscope, with a 63X objective (Plan Apochromat, oil immersion, NA=1.4).

### Fluorescence microscopy

The cells were plated on coverslips (12mm #1, ThermoFisher). THP-1 cells were differentiated with PMA (160 nM) for 48h and then infected transduced/infected (MOI and time post-infection indicated in figure legends). For late reverse transcripts or RNA visualization the cells were transduced with OR-GFP LV (MOI 0.5) or MCP-GFP LV (MOI 2) and 48 h later infected with HIV-1 ANCH3 or HIV-1 ANCH3 MS2, respectively. The medium was always supplemented with PMA. For HIV-1 ANCH3 imaging in HeLa P4R5, the cells were transduced with OR-GFP LV (MOI 0.2) and then infected 24 hours later with HIV-1 ANCH3 or HIV-1 IN_D116A_ ANCH3 + NEVIRAPINE (NEV) 10 µM or RALTEGRAVIR (RAL) 20 µM drugs were used to block respectively DNA synthesis or integration. The day of fixation the cells were washed with PBS and fixed with 4% PFA for 15 minutes. For protein staining, cells were treated with glycine 0.15% for 10 min, permeabilized with 0.5% Triton X-100 for 30 min and blocked with 1% bovine serum albumin (BSA) for 30 min. All antibody incubations were carried out at room temperature, in the dark, in humid chamber, 1h with primary antibodies and 45 min with secondary antibodies. Washes between antibody incubations and antibodies dilution were done in 1% BSA. Primary antibodies were diluted as follows: anti-HA 1:500 (Roche #11867423001), anti-CPSF6 1:400 (Novus Biologicals #NBP1-85676), anti-SC35 1:200 (Abcam #ab11826), anti-LEDGF 1:200 (BD Bioscience #611715). Secondary antibodies used were the following: Donkey anti-Rabbit Cy3 1:1000 (Jackson Lab #711-165-152), Goat anti-Rabbit Alexa-488 1:300 (Invitrogen #A32731), Goat anti-Mouse Alexa-647 1:300 (Invitrogen #A21235); Donkey anti-Rat Alexa-488 (Invitrogen # A21208) or Goat anti-Rat Alexa-647 (Invitrogen #A21247) 1:100 for IN-HA. For viral DNA detection via EdU incorporation, the cells were infected in presence of EdU (5 µM) for 3 days (adding new RPMI+EdU medium every day) and after fixation the click-chemistry was performed before immunostaining as in Rensen et al., 2021 (Rensen et al. 2021). Finally, cells were stained with Hoechst 33342 1:10000 (Invitrogen #H3570) for 5 minutes. Coverslips were mounted on glass slides (Star Frost) with Prolong Diamond Antifade Mountant (Life Technologies #P36970). Confocal microscopy was carried out with a Zeiss LSM700 inverted microscope, with a 63X objective (Plan Apochromat, oil immersion, NA=1.4).

### FRAP and time-lapse microscopy

For all time-lapse studies the cells were plated in a polymer-coverslip bottom µ-Dish 35 mm (ibidi #81156). Two million of THP-1 cells were differentiated for 48 h and then transduced with CPSF6 mNeonGreen LV (MOI 0.01) for 3 days. Next, the cells were infected with HIV-1 and the live imaging was performed from 24 to 72 h p.i. For CPSF6 clusters fusion studies CPSF6 mNeonGreen positive cells were imaged every 5 min in 2D with a Biostation IM-Q (Nikon).

For Fluorescence Recovery After Photobleaching (FRAP) experiments, the selected region of interest was irradiated for 200µs/pixel with 488 nm/561nm laser and after bleaching the frames were acquired every 5 to 10 seconds for 5 minutes through a Ti2E inverted microscope (Nikon), based on a CSU-W1 spinning-disk (Yokogawa), using a 60X objective (Plan Apochromat, oil immersion, NA=1.4).

Experiments of live tracking of IN-Ruby (GIR) and viral DNA (OR-GFP) were performed in differentiated THP-1 cells transduced with OR-GFP LV (MOI 5). Two days post-transduction the cells were infected with HIV-1 ANCH3 GIR virus (MOI 30). Different cells were imaged starting from 45 h p.i. to 96 h p.i., every 5 minutes in 3D (stacks spacing 0.3 µm) with a Ti2E inverted microscope (Nikon), based on a CSU-W1 spinning-disk (Yokogawa), using a 60X objective (Plan Apochromat, oil immersion, NA=1.4).

### Imaging Analysis and Statistics

All data were analyzed with GraphPad Prism 9 (GraphPad Software, La Jolla California USA, www.graphpad.com), statistic tests are indicated in figure legends. All images and videos were analysed in Fiji (Schindelin et al. 2012) and Icy software 2.2.1.0 (de Chaumont et al. 2012). In particular, CPSF6 clusters segmentation was performed though HK-Means block in Icy (Dufour et al. 2008) followed by ROI statistics exportation (e.g., Mean Intensity, Volume, Area, Sphericity/Circularity defined as the normalised ratio between the contour and interior of the ROI, expressed as a percentage (100% for a circle or sphere). For the vDNA/vRNA distance form SC35 or HIV-1 MLO the 3D confocal images were processed for multi-channel image splitting, vDNA and SC35 segmentation. The automated 3D segmentation of cell nuclei included a preliminary non-local means denoising step to cope with the strong signal heterogeneity and the segmentation of SC35 was readjusted to low contrast conditions. The 3D boundary-to-boundary distance was computed between each vDNA or vRNA spot and its closest Hoechst-marked nuclear envelope border, SC35 speckle border or the closest HIV-1 MLO (SC35+CPSF6) border.

## Acknowledgements

We wish to thank Marion Louveaux and Dmitry Ershov for image analysis help, Philippe Souque, Selen Ay, for experimental help. We thank Fabrizio Mammano, Edouard Bertrand, Florian Mueller, Roy Parker and Edward Campbell for sharing reagents. We gratefully acknowledge the UtechS Photonic BioImaging platform (Imagopole), C2RT at Institut Pasteur, supported by the French National Research Agency (France BioImaging; ANR-10–INSB–04; Investments for the Future). We thank the NIH AIDS Reagents program to support us with precious reagents. We thank Addgene for the reagents, Franck Gallardo CEO at the NeoVirTech, ActiveMotif for ChIP help. This work was funded by the ANRS REACTing grant ECTZ88162 with a nominative PhD student fellowship ECTZ88177 for V.S., the Sidaction/FRM grant VIH20170718001, the Pasteur Institute.

## Supplementary Material

### Supplementary Figures

**Supplementary Figure S1.**
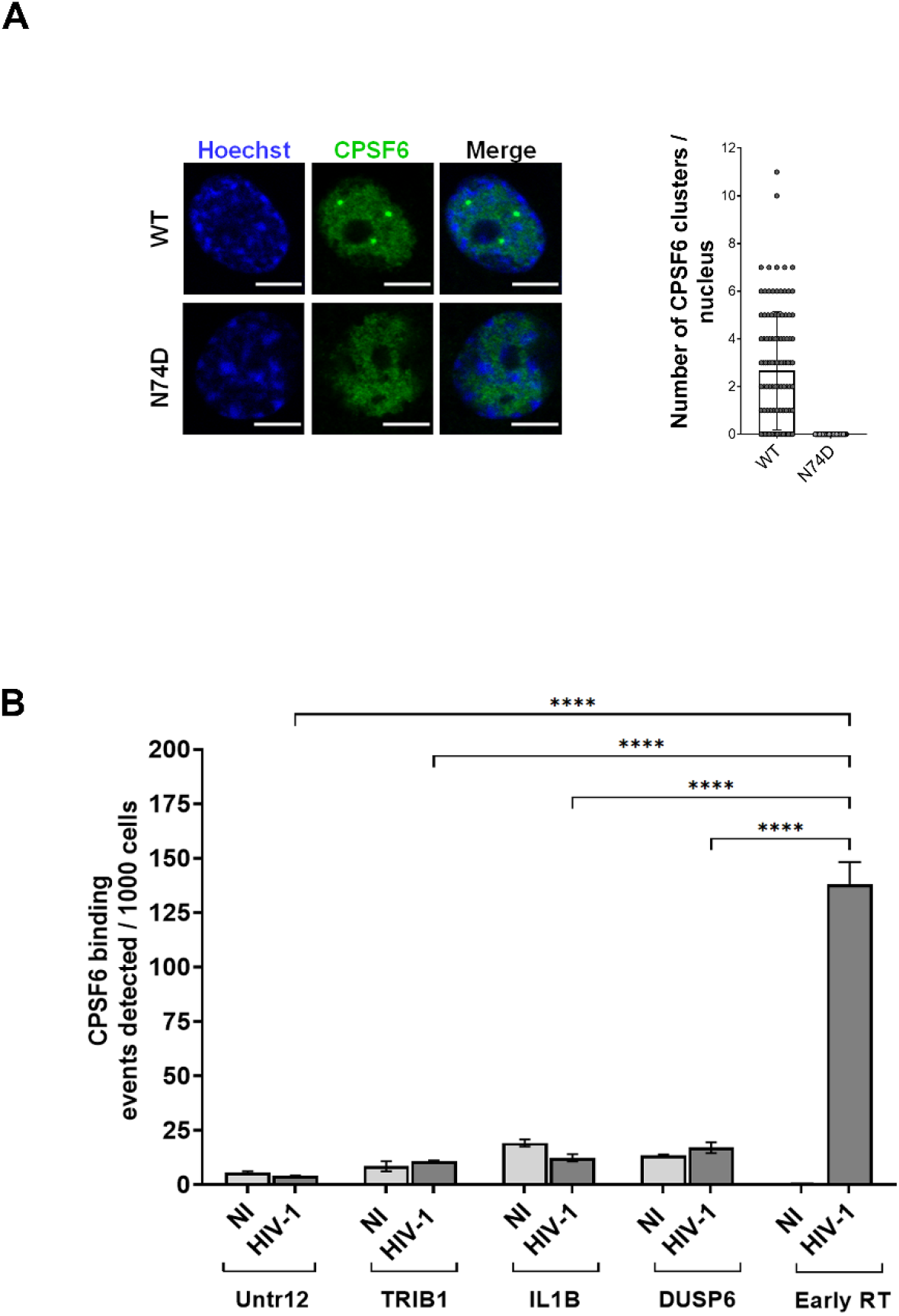
**A)** Confocal microscopy images of THP-1 cells infected with CA WT and CA N74D HIV-1 (MOI 5, 3 days p.i.). On the right, CPSF6 clusters count per cell ± SD (n cells = 108, 120). Unpaired t test, ****=p ≤ 0.0001. **B)** The histogram plot shows the number of copies ± SD of random genes in CPSF6 ChIP normalized for the input, compared to HIV-1 early reverse transcripts (RT) also normalized for the input (Figure 1C). THP-1 cells infected with HIV-1 (MOI 5), 2 days p.i. One-way ANOVA followed by Tukey’s multiple comparison test, ****=p ≤ 0.0001.

**Supplementary Figure S2.**
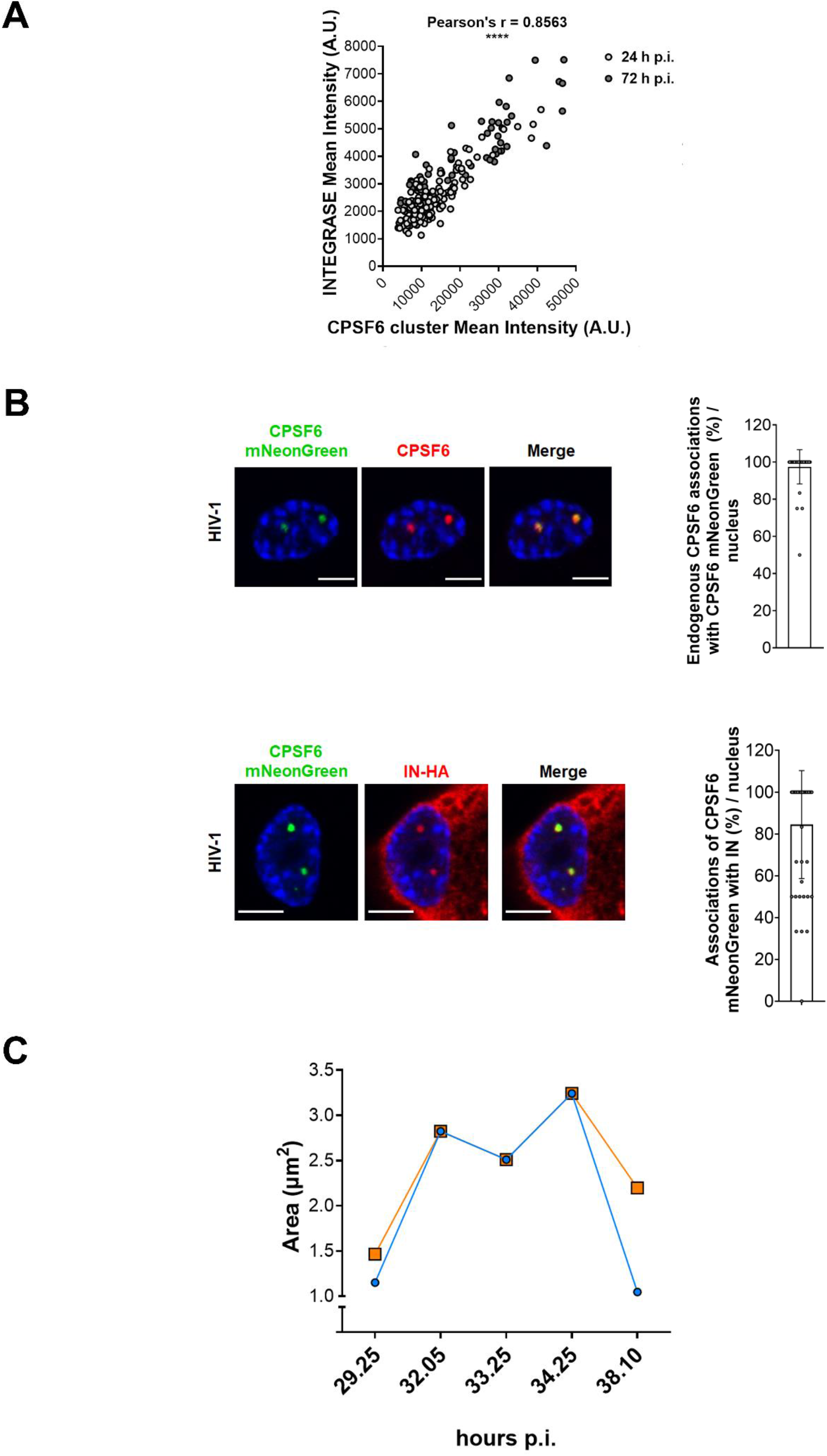
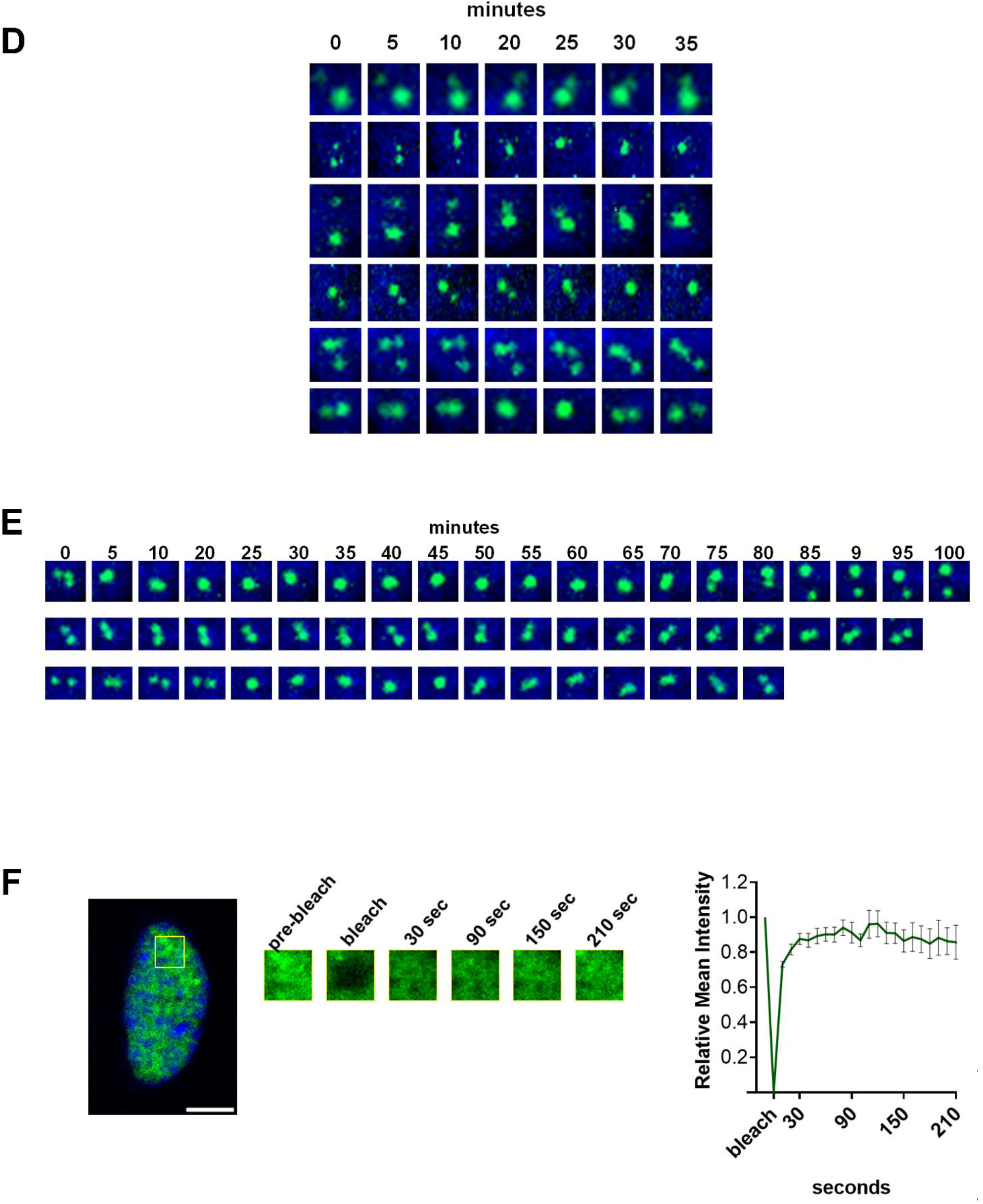
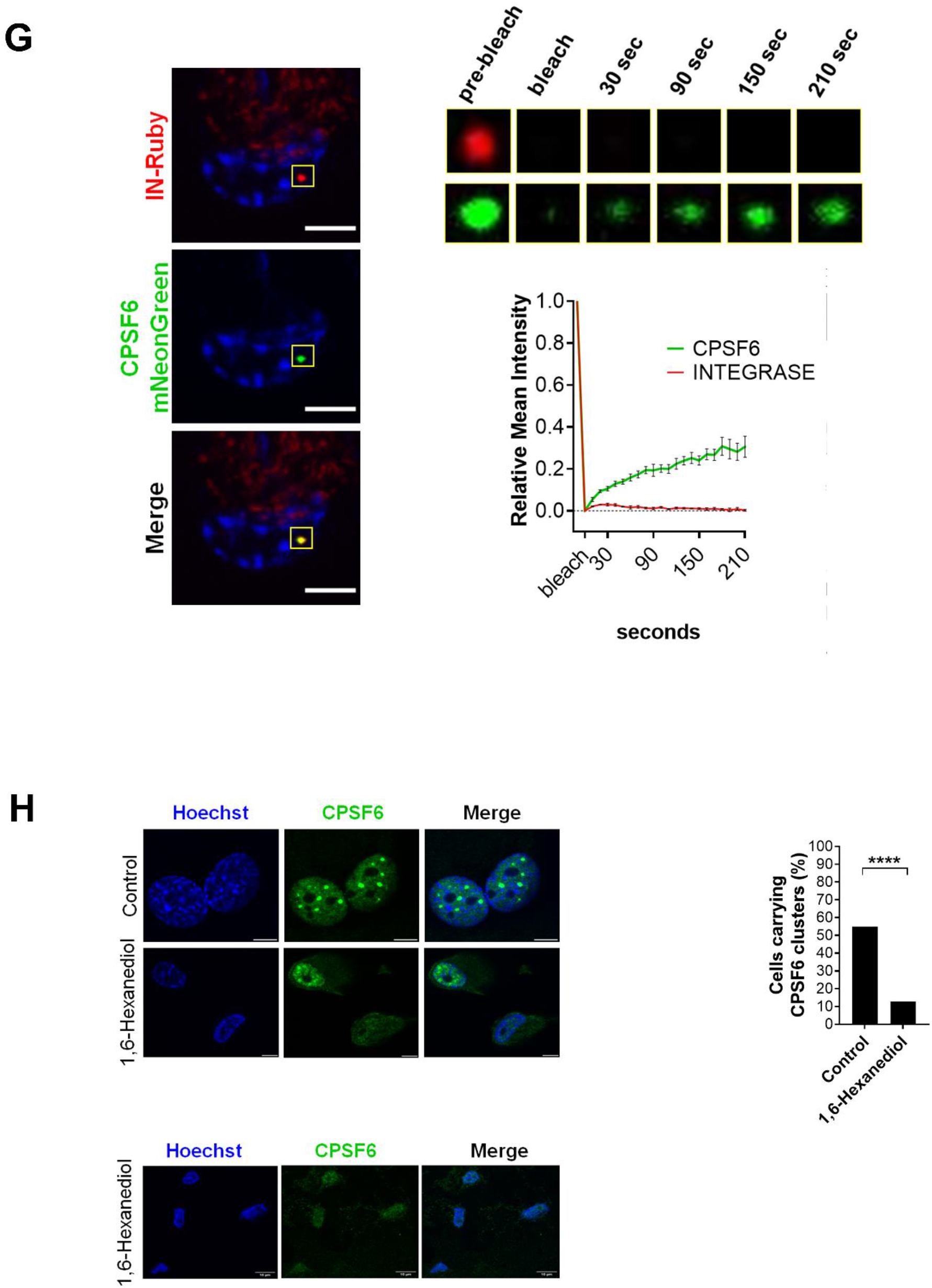
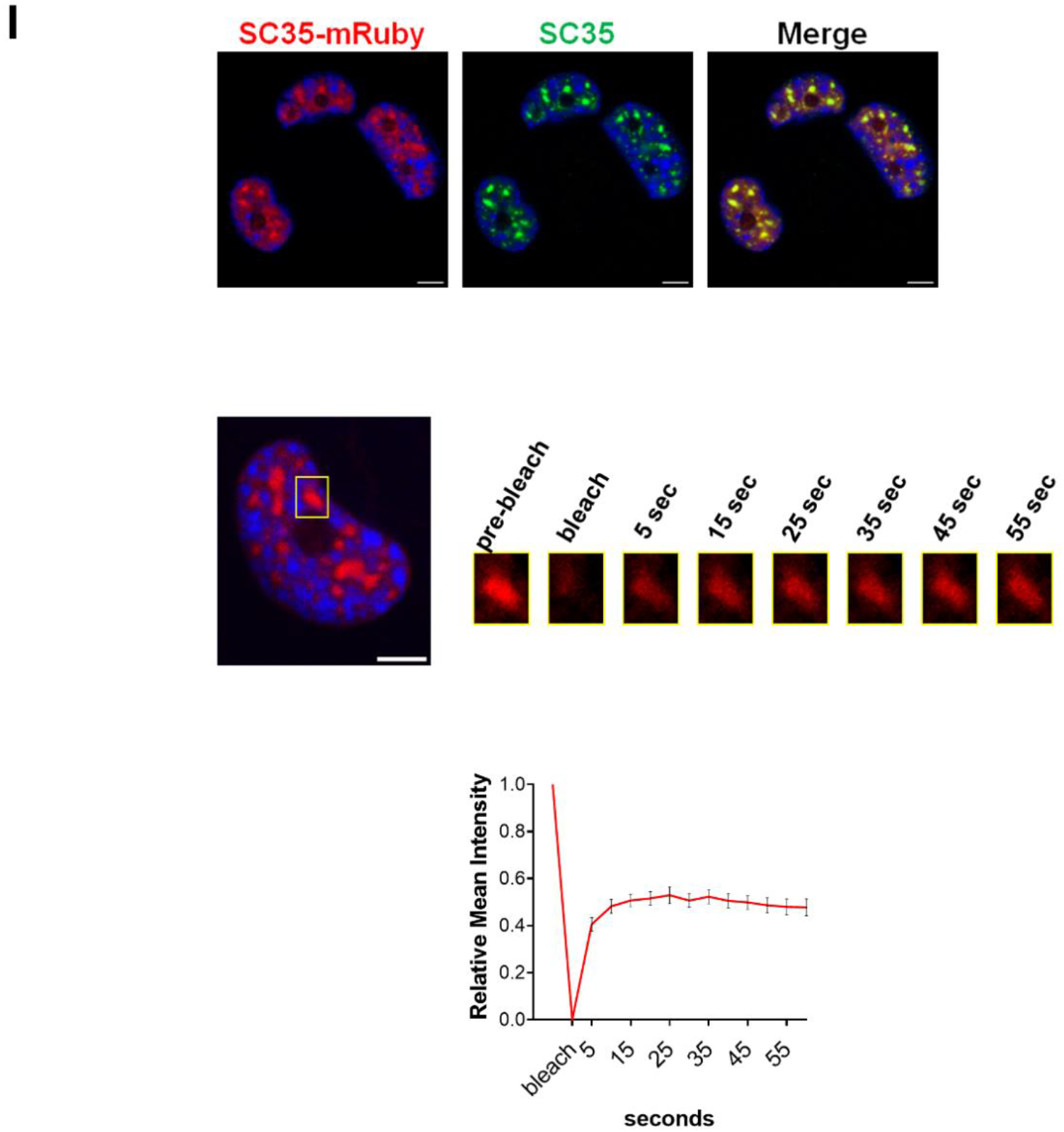
**A)** Pearson’s correlation of the mean intensity of CPSF6 clusters with the colocalizing IN focus (****=p ≤ 0.0001) in THP-1 cells infected with HIV-1 (MOI 5) + NEV (10 μM), 24 h and 72 h p.i. 3D analysis. **B)** Confocal microscopy images of THP-1 cells expressing CPSF6 mNeonGreen infected with HIV-1 (MOI 10, 3 days p.i.), stained with anti-CPSF6 antibody (red, above) or with anti-HA (INTEGRASE) (red, below). The graphs show the percentage ± SD of CPSF6 clusters labelled with the antibody that colocalizes with CPSF6 mNeonGreen clusters (n=45 cells), and the percentage ± SD of CPSF6 mNeonGreen clusters colocalizing with IN-HA foci (n=49 cells). **C)** The graph shows the area along the memebraneless organelles fusion-fission event (Figure 2C and Supplementary Video S1). **D)** Short-timing and **E)** Long-timing fusion/fission events from time-lapse microscopy in 2D of THP-1 cells expressing CPSF6 mNeonGreen infected with HIV-1 (MOI 10). Videos acquired 5 min/frame between 24 to 72 h p.i. **F)** Frames extracted form a FRAP time-lapse in uninfected THP-1 cells expressing CPSF6 mNeonGreen. The graph shows the recovery of the signal curve ± SEM (8 FRAP). Pre-bleach signal is set to 1 and bleach signal is set to 0. **G)** Frames extracted form a FRAP time-lapse in THP-1 expressing CPSF6 mNeonGreen infected with HIV-1 GIR (MOI 5, 3 days p.i.). Integrase focus (red) and CPSF6 cluster (green) were in turn bleached with the correspondent laser (561nm/488nm) the foci. The graph shows the recovery of the signal curves ± SEM (19 CPSF6 FRAP, 4 INTEGRASE FRAP). Pre-bleach signal is set to 1 and bleach signal is set to 0. **H)** Confocal microscopy images of THP-1 cells infected with HIV-1 (MOI 5). Twenty-four hours p.i. cells were treated with 10% 1,6-Hexanediol (1,6-Hexa) for 10 minutes. On the right, percentage of cells with CPSF6 clusters (n cells = 60, 55). Fisher’s exact test, ****=p ≤ 0.0001. **I)** On the top confocal microscopy images of THP-1 cells expressing SC35-mRuby (3 days post transduction) stained with anti-SC35 antibody. On the bottom frames extracted form a FRAP time-lapse in uninfected THP-1 cells expressing SC35-mRuby. The graph shows the recovery of the signal curve ± SEM (14 FRAP). Pre-bleach signal is set to 1 and bleach signal is set to 0. Scale bars: 5 μm, unless indicated.

**Supplementary Figure S3.**
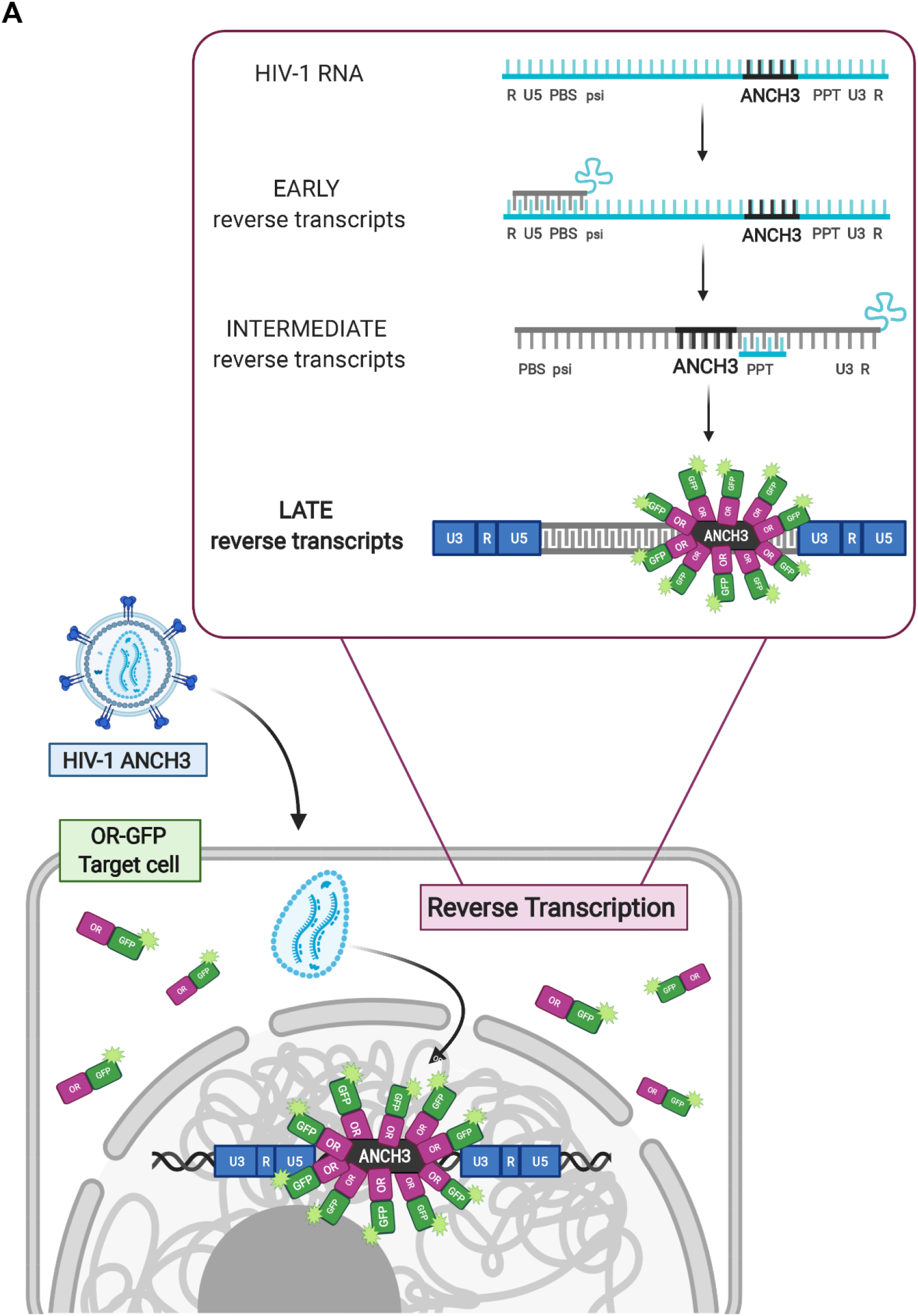
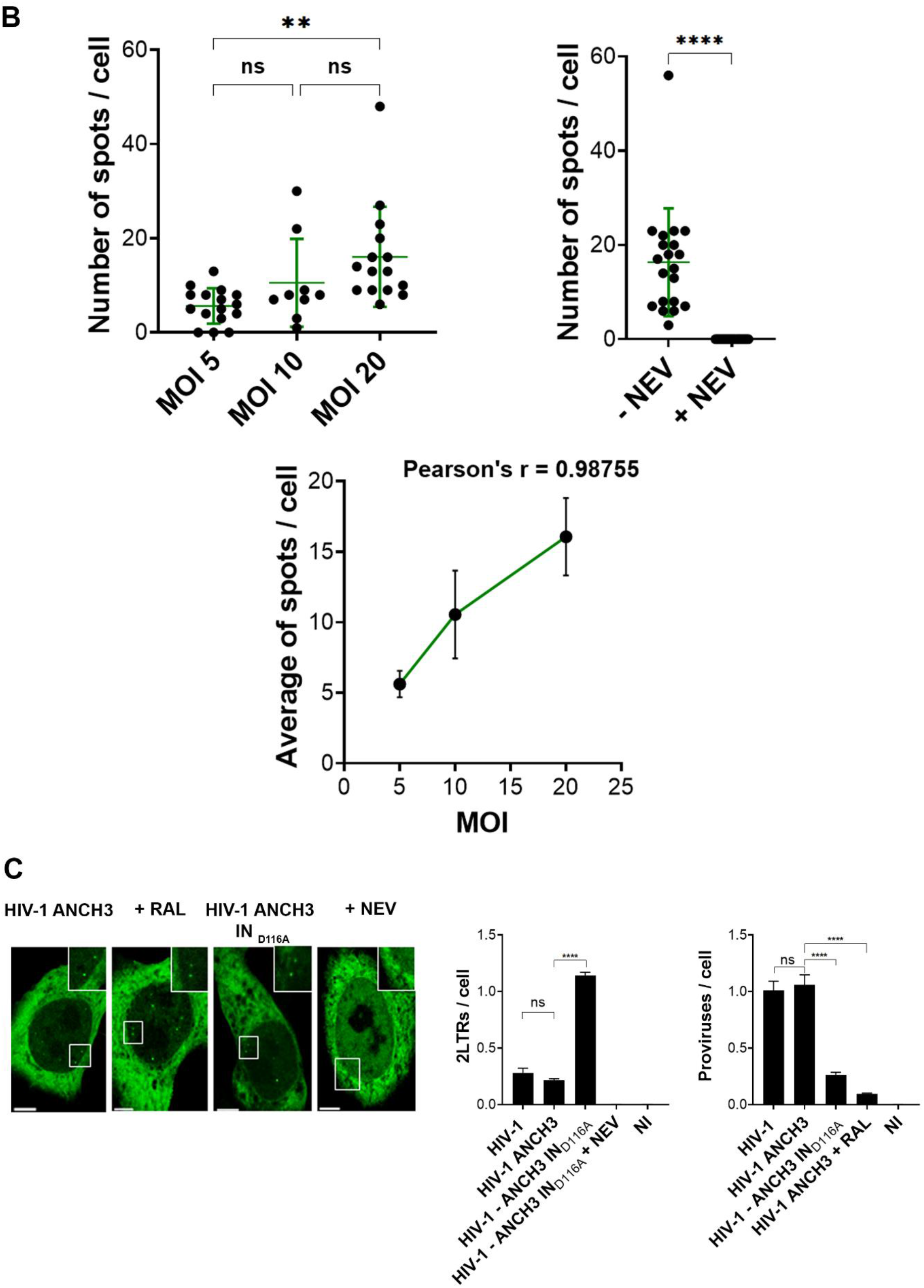
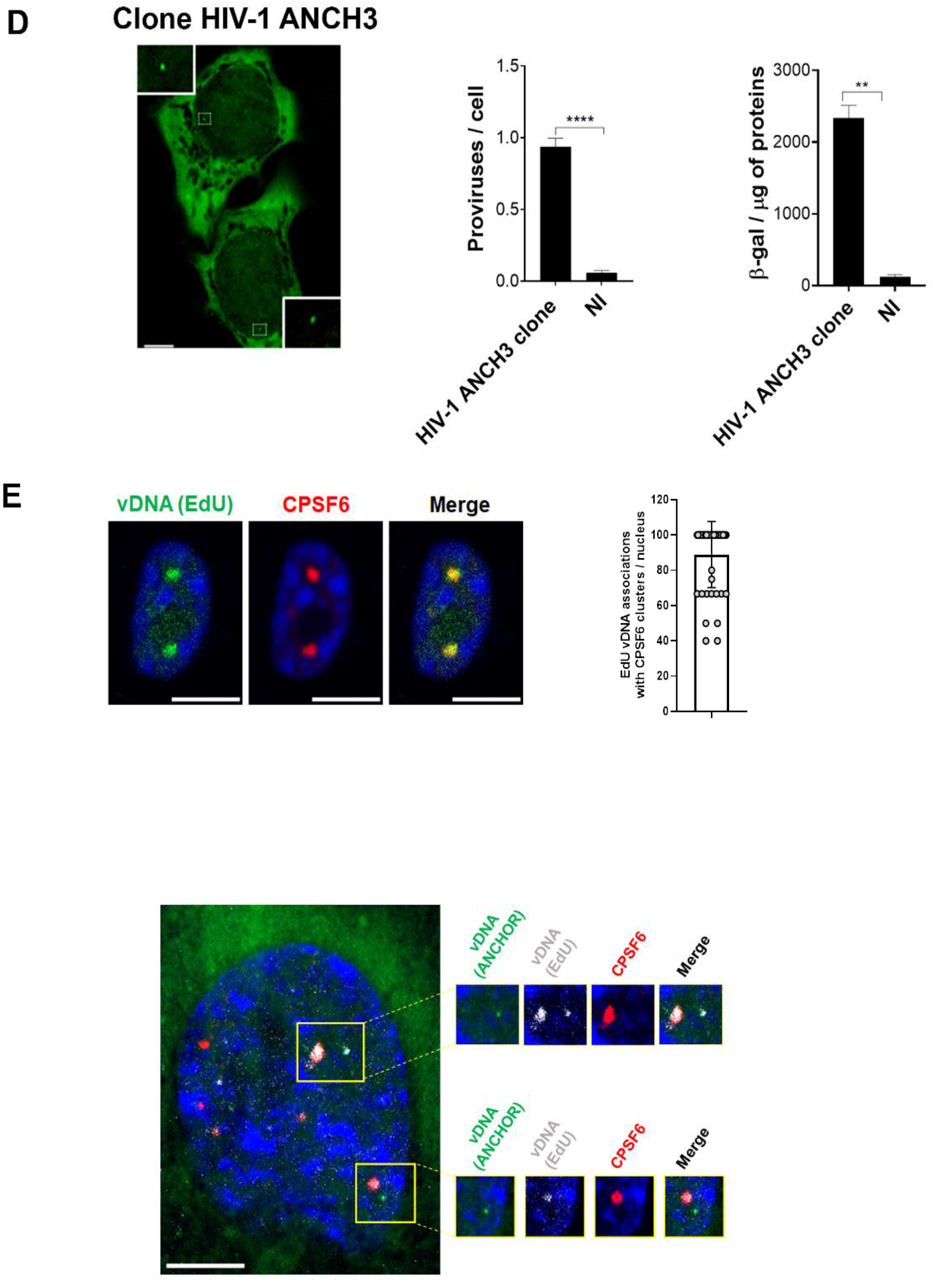
**A)** Scheme of HIV-1 ANCHOR system created with BioRender.com. ANCH3 sequence has been cloned in the HIV-1 genome in place of Nef gene (HIV-1 ANCH3). The late retrotranscribed DNA of HIV-1 ANCH3 can be visualized in cells stably expressing the OR-GFP protein, thanks to OR accumulation on the double-stranded ANCH3 sequence. Viral DNA detection occurs mostly in the nucleus, where late reverse transcripts are abundant and are more likely to be exposed to the binding of OR-GFP. **B)** Analysis in HeLa P4R5 cells expressing OR-GFP infected with HIV-1 ANCH3, 24 h p.i. The plots show: the vDNA spots count per cell for each MOI used and the comparison between the count in infected cells treated or not with NEV (10μM). Pearson’s correlation of the average spots count with the increasing MOI used (n cells = 16, 9, 15) ± SEM. **C)** Confocal images of HeLa P4R5 cells expressing OR-GFP infected with HIV-1 ANCH3, HIV-1 ANCH3 + RAL (20 μM), HIV-1 ANCH3 IND116A and HIV-1 ANCH3 + NEV (10 μM) (MOI 30, 24 h p.i.). The histogram plots of 2-LTR circles and ALU-PCR represent the post-nuclear formation of episomal form and integration rates, respectively. One-way ANOVA followed by Tukey’s multiple comparison test, ns=not-significant, ****=p ≤ 0.0001. **D)** Confocal image of HIV-1 ANCH3 provirus in an infected clone. On the right, histogram plot of ALU-PCR and β-galactosidase expression in the infected clone compared to uninfected cells (NI). Unpaired t test, **=p ≤ 0.01, ****=p ≤ 0.0001. **E)** Confocal microscopy images of THP-1 cells infected with HIV-1 in presence of EdU (MOI 20, 3 days p.i.) and quantification of EdU foci associated with CPSF6 clusters per nucleus (n=45 cells). The image below shows a confocal microscopy image of THP-1 infected with HIV-1 ANCH3 in presence of EdU (MOI 20, 3 days p.i.). Scale bars: 5 μm.

**Supplementary Figure S4.**
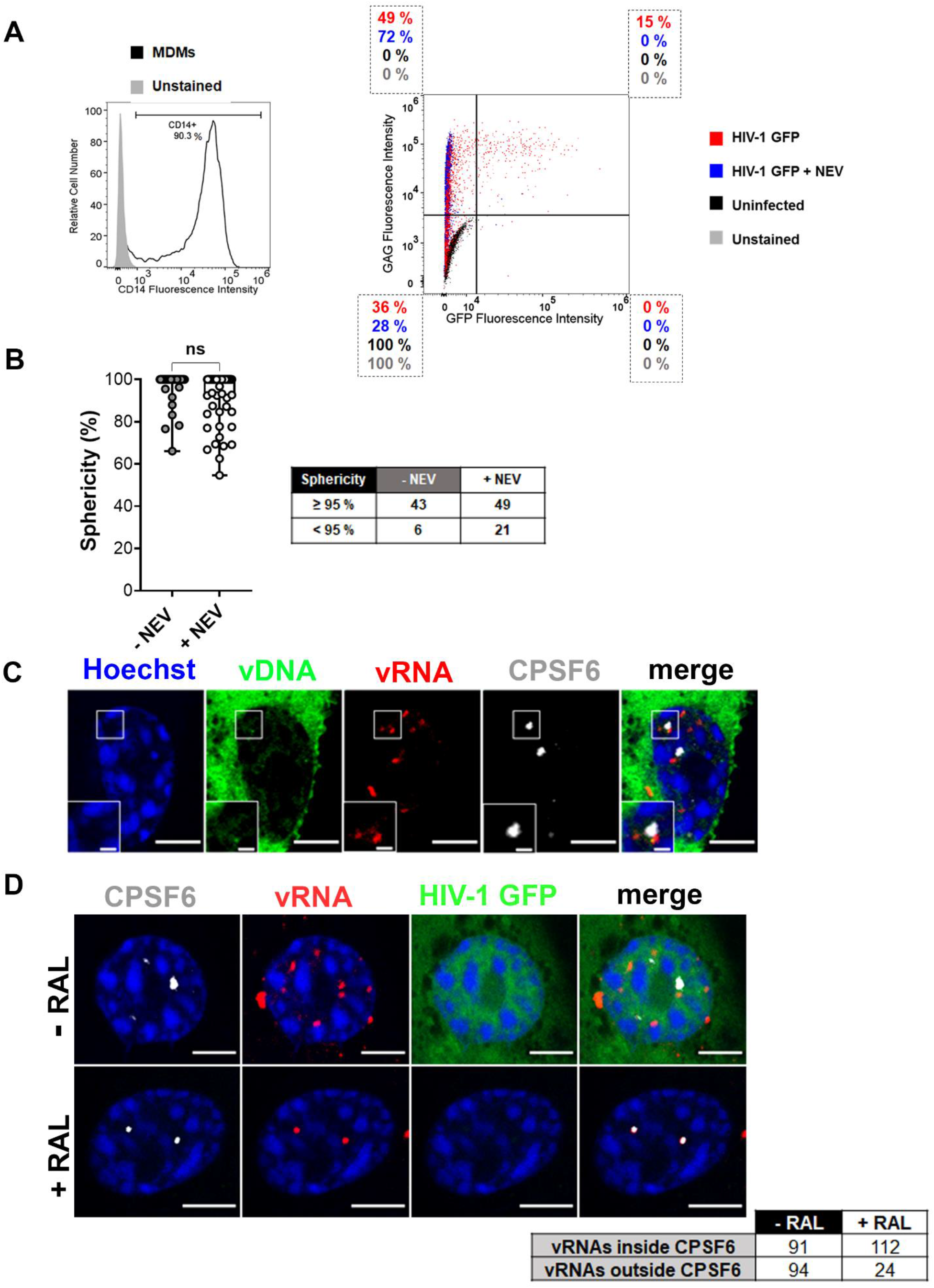
**A)** The FACS histogram displays CD14+ expression of the MDMs in this study. The FACS dot plot shows the correlation of GAG signal and GFP expression, with the related percentages per population (MOI 20, 2-3 days p.i.). **B)** Box plot of the sphericity of CPSF6 clusters in infected MDMs. Unpaired t test, ns=not-significant. **C)** Confocal image of Immuno-RNA FISH in THP-1 cell expressing OR-GFP infected with HIV-1 ANCH3 (MOI 20, 3 days p.i.). **D)** Confocal images of Immuno-RNA FISH in THP-1 cells infected with HIV-1 GFP ± RAL (20 μM) (MOI 10, 2 days p.i.). Scale bar: 5 μm, inset: 1 μm.

**Supplementary Figure S5.**
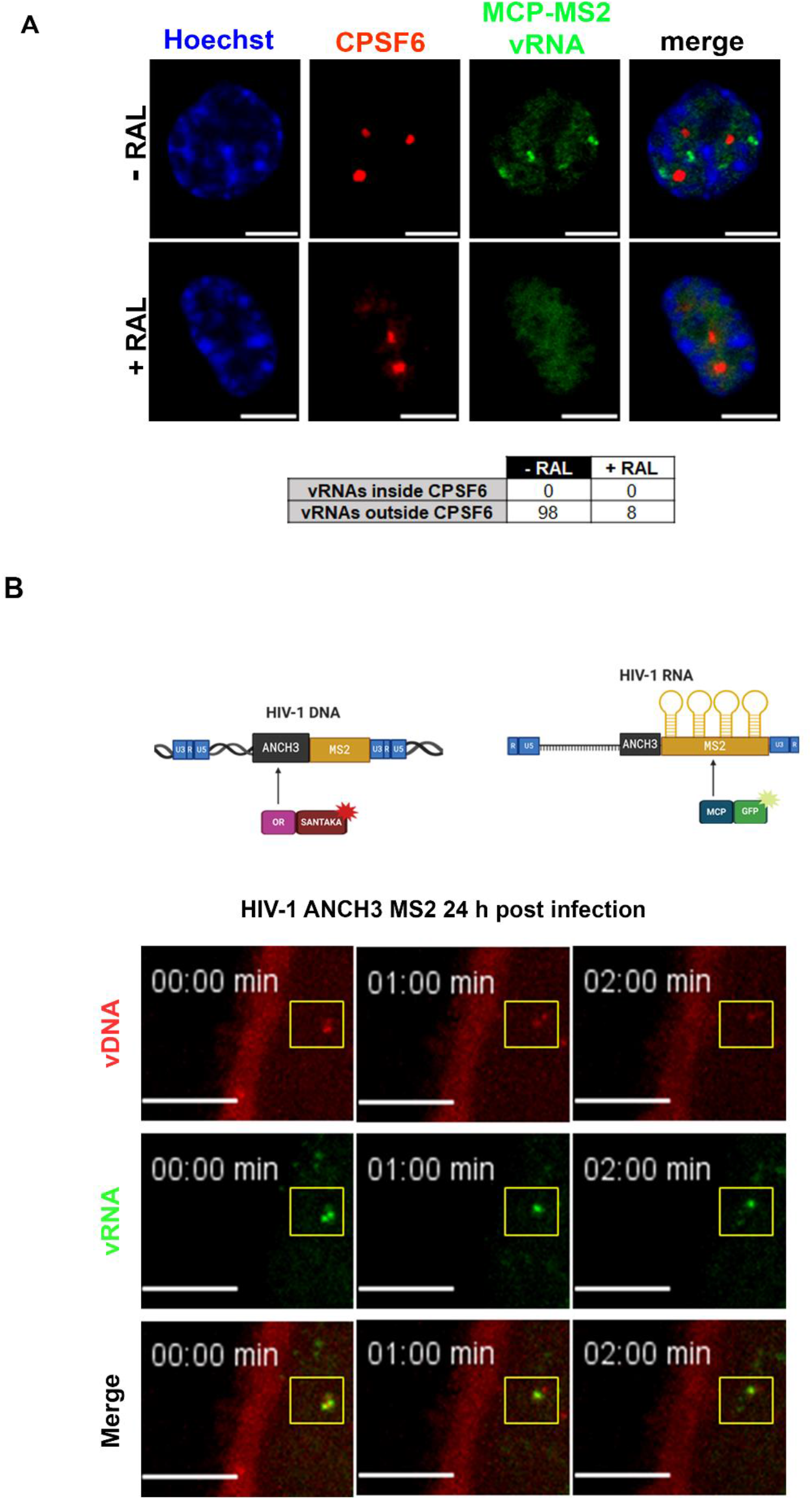
**A)** Confocal images of THP-1 cells expressing MCP-GFP infected with HIV-1 ANCH3 MS2 ± RAL (20μM), (MOI 2.5, 3 days p.i.). **B)** The cartoon model shows the double labelling system: the HIV-1 genome is tagged both with ANCH3 and MS2 sequences. The differently fluorescent reporters allow the simultaneous detection of vDNA through the binding of OR-GFP to ANCH3 sequence and of vRNA through the binding of MCP to MS2 stem loops. Created with BioRender.com. Frames from a time-lapse microscopy of HeLa MCP-GFP cells transduced with OR-SANTAKA LV (MOI 1) and 48 hours after infected with HIV-1 ANCH3 MS2 (MOI 50), 24 h p.i.. Projection of 6 Z slices, spacing 0.2 μm with Ti2E inverted microscope (Nikon), based on a CSU-W1 spinning-disk (Yokogawa), using a 60X objective (Plan Apochromat, oil immersion, NA=1.4). Scale bar: 5 μm.

**Supplementary Figure S6.**
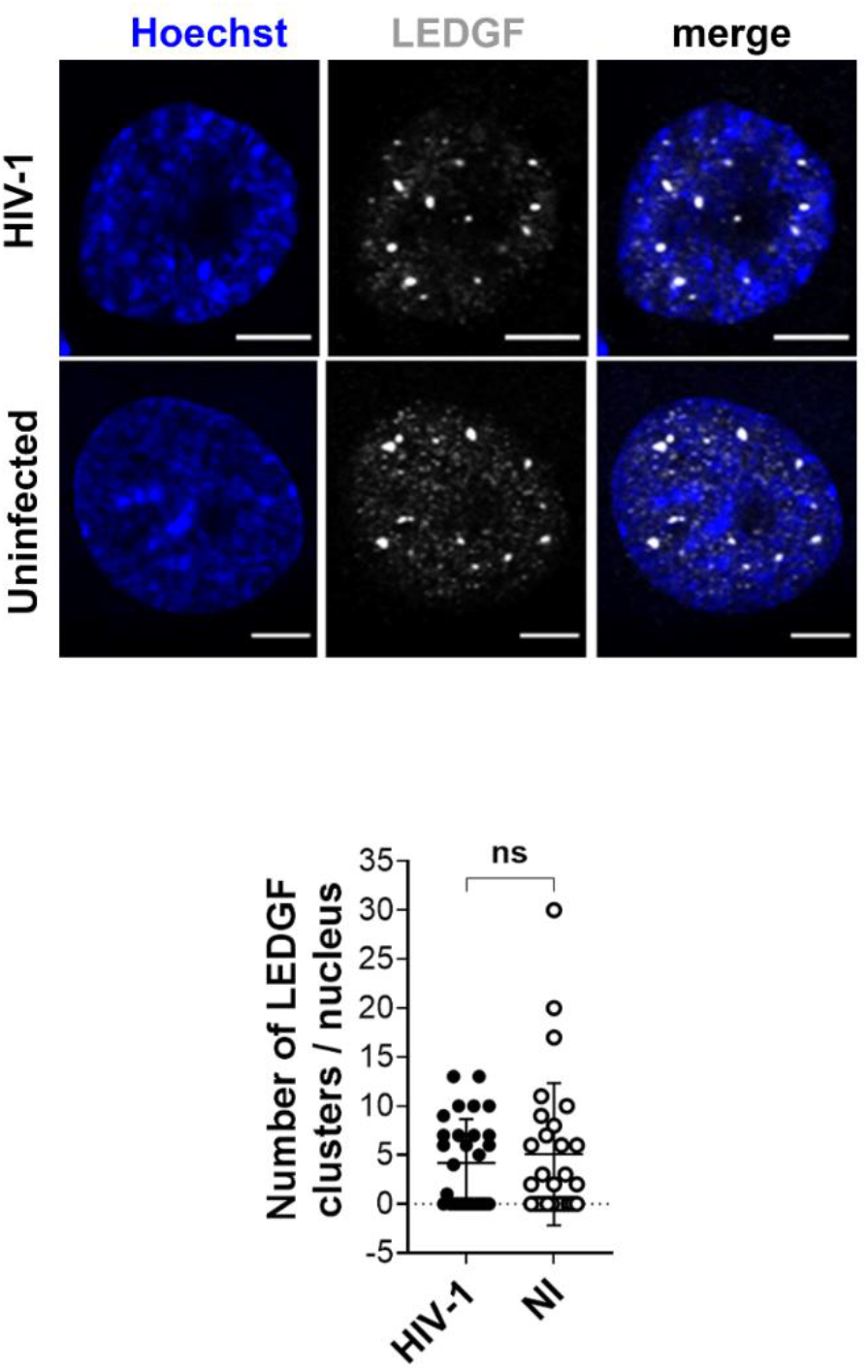
**A)** Confocal images of THP-1 cells infected with HIV-1 (MOI10, 3 days p.i.) compared to uninfected cells. The scatter plot shows the count of LEDGF clusters per nucleus (n = 29 cells (HIV-1), 28 cells (NI)); unpaired t test, ns = not-significant, Scale bar: 5 μm.

### Supplementary Video Legends

**Supplementary Video S1. CPSF6 clusters dynamics in HIV-1 infected cells.** Time-lapse microscopy of THP-1 cells expressing CPSF6 mNeonGreen infected with HIV-1 (MOI 10). Time post infection is indicated on the top. It is possible to appreciate different fusion/fission events of CPSF6 clusters (green). Representative of 6 independent experiments. Scale bar: 5 μm.

**Supplementary Video S2. FRAP time lapse of CPSF6 clusters in HIV-1 infected cells.** FRAP time-lapse in THP-1 cells expressing CPSF6 mNeonGreen infected with HIV-1 (MOI 10, 3 days p.i.). The first 3 frames are pre-bleaching, frame 4 is the bleach and from frame 5 it starts the recovery. Representative of 5 independent experiments. Scale bar: 5 μm.

**Supplementary Video S3. Generation and separation of ds vDNA from the IN focus in live.** Time-lapse microscopy in THP-1 cells infected with HIV-1 ANCH3 GIR virus (MOI 30, 80 h p.i.). The vDNA (green) signal generates from the integrase (IN) proteins focus (red). The video is a montage of continuous 2D frames (5 min/frame). Scale bar: 5 μm.

### Supplementary Methods

**Supplementary Table S1.**
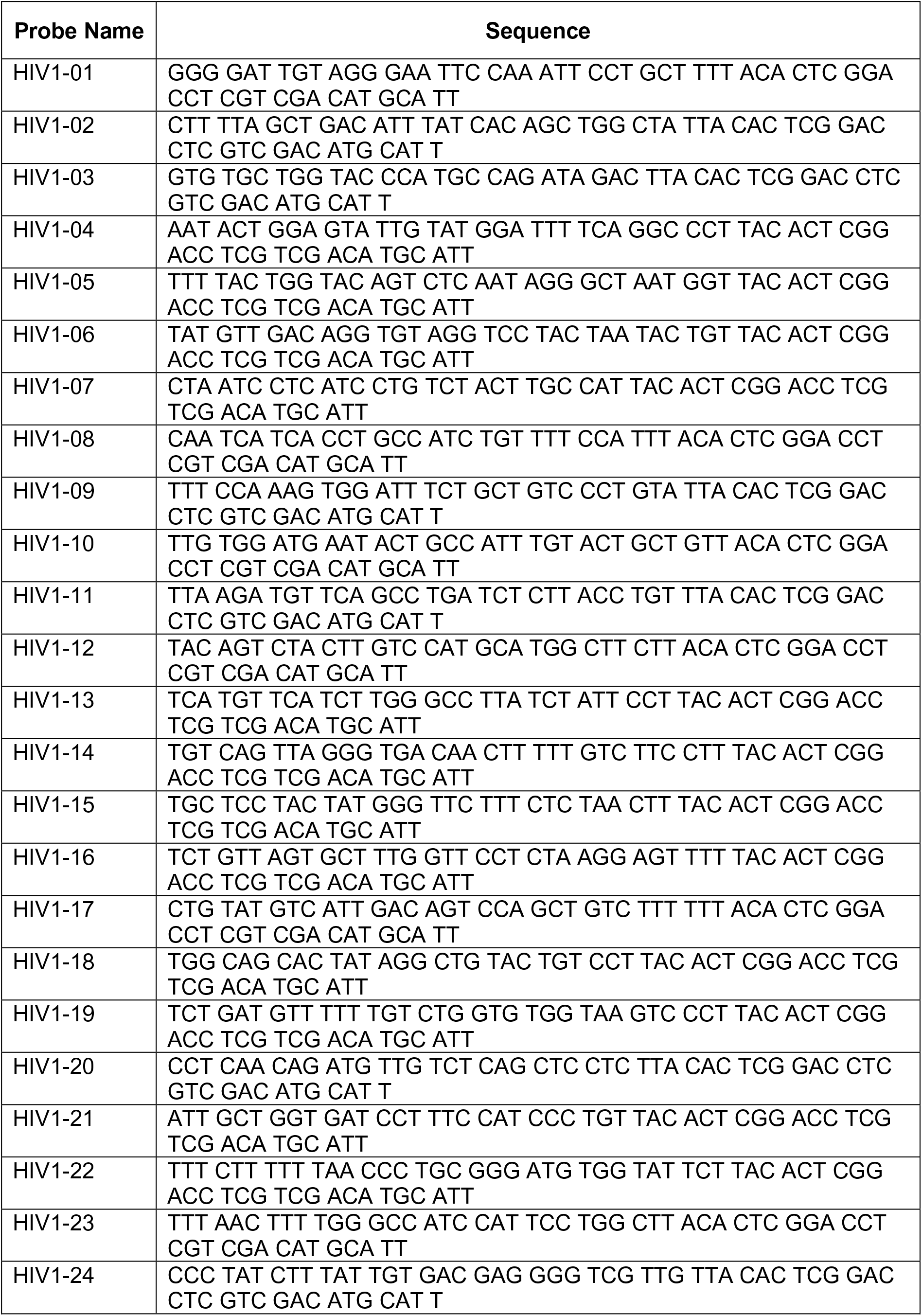
Probes used for RNA FISH against pol.

#### Quantitative PCR and primers

For HIV-1 ANCH3 DNA forms quantification in HeLa P4R5 cells, a total DNA of 10^6^ infected cells (MOI 30) was extracted at 6 hours post infection for Late Reverse Transcripts (LRTs) qPCR and 24 hours post infection for 2LTR-circles and ALU-PCR, through QIAmp DNA micro kit (QIAGEN #56304). Real-Time PCR of LRTs was performed to assess DNA synthesis and used to normalize qPCR data on viral input, the reactions were carried on in 20 µL, in iTaqUniversal SYBR Green Supermix (Bio-Rad #1725124) using primers for U3 sequence: U3 FX: 5’- TTCCGCTGGGGACTTTCCAGGG-3’, U3 RX: 5’-AGGCTCAGATCTGGTCTAACC-3’. Real-Time PCR of 2LTR-circles was used to assess nuclear import efficiency. Reactions were performed in 20 µL, in Maxima Probe/ROX qPCR Mastermix (ThermoFisher #K0232) using primers for 2LTR-circle junction: 2LTR FX: 5’-AACTAGGGAACCCACTGCTTAAG-3’, 2LTR RX: 5′-TCCACAGATCAAGGATATCTTGTC-3′, 2-LTR probe: 5’-(FAM)-ACACTACTTGAAGCACTCAAG-GCAAGCTTT-(TAMRA)-3’. Proviral integration quantification of HeLa P4R5 infected cells and HIV-1 ANCH3 clone was performed through ALU PCR, consisting in a first non-kinetic PCR step in 50 μL using Platinum SuperFi DNA Polymerase kit (ThermoFisher #12351250) for the amplification of ALU-U3 fragments and in a second step of HIV-1 specific qPCR reaction in 20 μL in Maxima Probe/ROX qPCR Mastermix (ThermoFisher #K0232). Primer Alu 166: 5’-TCCCAGCTACTCGGGAGGCTGAGG-3’, Alu 2: 5’-GCCTCCCAAAGTGCTGGGATTACAG-3’, LambdaU3: 5′-ATGCCACGTAAGCGAAACTTTCCGCTGGGGACTTTCCAGGG-3′ for ALU PCR. Lambda: 5’-ATGCCACGTAAGCGAAACT-3’, U5: 5’-CTGACTAAAAGGGTCTGAGG-3’, Probe: 5’-(FAM)-TTAAGCCTCAATAAAGCTTGCCTTGAGTGC-(TAMRA) for qPCR. In all experiments β-actin detection was used for normalization. β-actin FX: 5’-AACACCCCAGCCATGTACGT-3’, β-actin RX: 5-CGGTGAGGATCTTCATGAGGTAGT-3’, β-actin probe: (FAM)-CCAGCCAGGTCCAGACGCAGGA-(BHQ1).

